# A one-dimensional individual-based mechanical model of cell movement in heterogeneous tissues and its coarse-grained approximation

**DOI:** 10.1101/485276

**Authors:** R. J. Murphy, P. R. Buenzli, R. E. Baker, M. J. Simpson

## Abstract

Mechanical heterogeneity in biological tissues, in particular stiffness, can be used to distinguish between healthy and diseased states. However, it is often difficult to explore relationships between cellular-level properties and tissue-level outcomes when biological experiments are performed at a single scale only. To overcome this difficulty we develop a multi-scale mathematical model which provides a clear framework to explore these connections across biological scales. Starting with an individual-based mechanical model of cell movement, we subsequently derive a novel coarse-grained system of partial differential equations governing the evolution of the cell density due to heterogeneous cellular properties. We demonstrate that solutions of the individual-based model converge to numerical solutions of the coarse-grained model, for both slowly-varying-in-space and rapidly-varying-in-space cellular properties. We discuss applications of the model, such as determining relative cellular-level properties and an interpretation of data from a breast cancer detection experiment.

## 1 Introduction

Biological tissues are heterogeneous and multi-scale by their very nature (Figure 1(*a*)). This heterogeneity exists at all scales from sub-cellular to cellular, and from cellular to tissue levels [1–3]. We focus on cellular interactions driven by mechanical stiffness which is of great importance in a variety of applications including epithelial tissue mechanics, cancer progression [4], cancer invasion and metastasis [5], stiffness as a biomarker in cancer detection [6–9], wound healing [10], and morphogenesis [11]. Tissue-level stiffness information [12] has been available for much longer than cellular-level stiffness data which requires advanced technology, such as atomic force microscopy [13–16]. However, difficulties arise in relating cellular-level data with tissue-level information when experiments are conducted and analysed at a single scale only. Mathematical modelling with *in silico* simulations provides a clear framework to explore these connections across biological scales.

**Figure 1:**
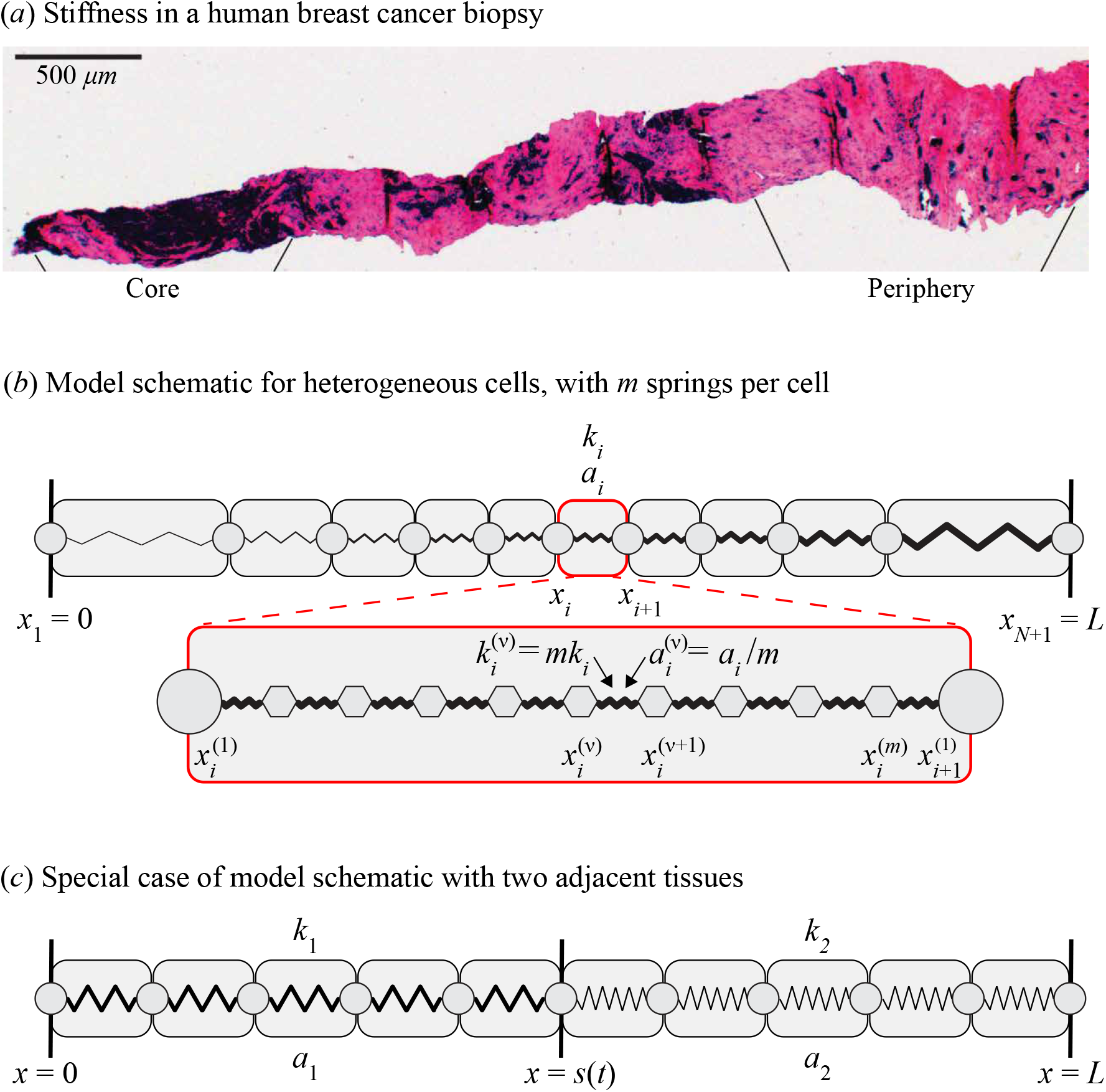
Stiffness heterogeneity in biological tissues. (*a*) Post atomic force microscopy histological overview of an entire breast cancer biopsy, where dark regions and pink regions are associated with low and high cell stiffness, respectively. Reproduced from [6] with permission. (*b*) Individual-based model schematic for arbitrarily heterogeneous tissue with *N* cells and *m* identical springs per cell. Cell *i* occupies the region *x*_*i*_(*t*) < *x* < *x*_*i*+1_(*t*) has cell stiffness *k*_*i*_ and resting cell length *a*_*i*_. Spring *ν* in cell *i*, occupies the region 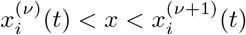, is prescribed with spring stiffness 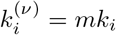 and resting spring length 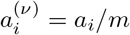. The first and final spring boundaries in cell *i* coincide with the cell boundary positions so that 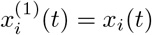 and 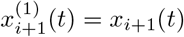 for all time. The cell and spring boundaries are shown as discs and hexagons, respectively. (*c*) Individual-based model schematic for a special case with two adjacent tissues, similarly this could model a heterogeneous tissue with two cell types. Cells in tissue *i* are prescribed with cell stiffness *k*_*i*_ and resting cell length *a*_*i*_ for *i* = 1, 2. Here each cell is represented with a single spring. The position of the interface between the two tissues is at *x* = *s*(*t*).

Mathematical models of cell populations are broadly classified as either discrete or continuum. Discrete models, reviewed in [17,18], include cellular automata models, cellular Potts models, cell-centre models [17], vertex models, subcellular-element models [19], and tensegrity models [20]. Discrete models explicitly describe cellular-level interactions but often lack macroscopic intuition. Continuum models on the other hand often provide no cellular-level information [21] but can be more adept at including concepts of macroscopic stiffness [22,23] and, for large numbers of cells, as in epithelial tissues, tend to be less computationally expensive. Hybrid intermediate models also exist which consider the multi-scale nature of the problem [24–26]. A range of models specifically examine the role of mechanics [27,28]. However, in this work we focus on models which relate cellular-level details to tissue-level outcomes. These models have been developed with a variety of coarse-graining techniques and assumptions, including the use of slowly varying and periodic assumptions on the heterogeneity in the model [29,30], correlation functions [31,32], and interaction forces from potentials [33]. Few of these models explore the role of stiffness. The work of Murray et al. [34–37] explicitly incorporates cell stiffness; they derive a nonlinear diffusion equation governing the evolution of the cell density in space and time, however the framework focuses exclusively on homogeneous cell populations. Here, we extend this framework to heterogeneous cell populations.

The key focus of this work is to present a novel coarse-grained system of partial differential equations governing the evolution of the cell density, cell stiffness and resting cell length, from a heterogeneous cell-based model of epithelial tissue mechanics. The cell stiffness and resting cell length are constant for each cell and are simply transported in space by cell movements. The motion in this model is governed by cell-cell interaction forces modelled with Hooke’s law. In extending the work of Murray et al. [34], we provide a more general derivation of the governing equations, see Section 2, which is robust to the inclusion of both slowly-varying-in-space and rapidly-varying-in-space cellular properties, see Section 3. We show that solutions from the discrete model converge to the corresponding continuum model solution, under appropriate scalings. Additional results in Section 3 show the model can be applied to interpret experimental and clinical observations relating to breast cancer detection. Key algorithms used to generate results are available on GitHub.

## 2 Model description

In this section we describe the individual-based model, which we refer to throughout this work as the discrete model, and derive a corresponding coarse-grained approximation in the form of system of partial differential equations, which we refer to as the continuum description. The continuum limit usually assumes that the number of discrete entities that makes up the system tends to infinity [29,38], while the size of the domain also tends to infinity, as in the thermodynamic limit, or the size of a length scale tends to zero, both in such a way that the ratio of the size of length scale to the number of discrete entities is fixed. Here, to maintain a fixed total tissue length and a fixed total number of cells in the continuum limit, we instead assume that each cell is internally represented by several identical springs. We then take the continuum limit by considering that the number of springs per cell tends to infinity whilst the spring length tends to zero.

### 2.1 Discrete model

In this work, the discrete model describes an epithelial tissue formed by cells in contact with each other. For simplicity, we assume that the tissue can be modelled as a one-dimensional chain of *N* cells with fixed total length *L*. Tissues in the body commonly evolve in confined spaces, for example imposed by bone tissues, and are subjected to strong geometric constraints so we fix the left tissue boundary at *x* = 0 and the right tissue boundary at *x* = *L*. This also allows us to focus on internal cellular heterogeneity. Alternate free boundary conditions are possible [34,39,40] but we do not focus on such free boundary conditions in this work. Each cell can have distinct mechanical properties (Figure 1). This model could be used to represent a single tissue with intrinsic heterogeneity or multiple adjacent tissues with different properties. Each cell interacts with its neighbour through an effective interaction force which could represent cell-cell adhesion [41] or compressive stresses [42]. We consider cell *i*, for *i* = 1, 2, …, *N*, to have its left boundary at *x*_*i*_(*t*) and its right boundary at *x*_*i*+1_(*t*), with *x*_1_(*t*) = 0 and *x*_*N*+1_(*t*) = *L* at all times. The cell has a prescribed cell stiffness, *k*_*i*_, and resting cell length, *a*_*i*_. Inside the tissue, Newton’s second law of motion governs the motion of each cell boundary such that

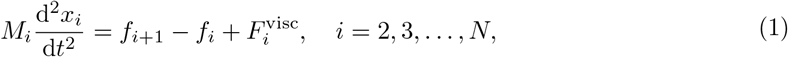

where *M*_*i*_ is the mass associated with cell boundary *i*, 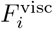 is the viscous force associated with cell boundary *i*, and we model interaction forces at cell boundary *i* using Hooke’s law,

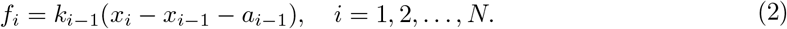

The viscous force experienced by cells, due to cell-medium and cell-matrix interactions, is modelled with 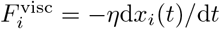, where *η* > 0 is the viscosity coefficient. Cells migrate in dissipative environments and this is commonly modelled by assuming that the motion is overdamped [34,43], hence the term on the left of Equation (1) is zero, giving,

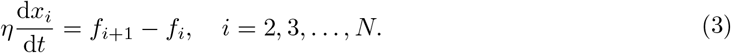

This model, as presented thus far, considers each cell to be represented by a single spring [34,40] which is sufficient to describe the discrete model. However, when we derive the continuum model, to maintain *L* and *N*, we represent each cell internally with *m* identical springs and we will later consider *m* ≫ 1, which corresponds to the spring length tending to zero, see Section 22.2. The corresponding discrete model for *m* springs per cell is now described. Cell *i* with boundaries *x*_*i*_ and *x*_*i*+1_ now has *m* + 1 spring boundaries, 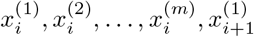, with 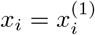 and 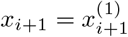, (Figure 1(*c*)). The cell length is related to the spring length through the scaling 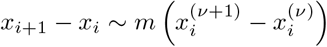 as *m* → ∞, and with equality for all *m* as *t* → ∞. Each spring *ν* in cell *i* is prescribed with a spring stiffness 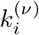 and resting spring length 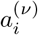 related to cell properties *k*_*i*_ and *a*_*i*_ through

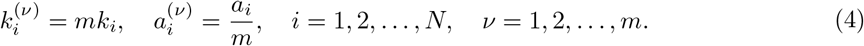

The viscosity coefficient for a cell boundary, *η*, is related to the viscosity coefficient for a spring boundary, *η*^(*ν*)^, through *η*^(*ν*)^ = *η/m*. Then the corresponding spring boundary equations are

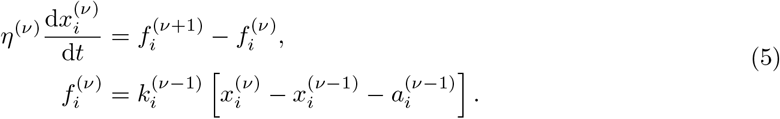

The scalings in Equation (4) and for the viscosity coefficient are chosen such that the cell boundary velocities are maintained and are independent of *m*, i.e. such that 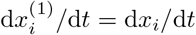. These scalings are supported by results from the discrete model with varying *m*, see Section 3.

The discrete model is governed by Equation (3) with the fixed boundary conditions for a system with a single spring per cell, and by Equation (5) with fixed boundary conditions for a system with *m* springs per cell. In each situation the discrete model forms a deterministic coupled system of ordinary differential equations that we can solve numerically, see Supplementary Material Section 1. We can also solve each system with an eigenmode decomposition to conveniently determine the long-time steady state solution.

### 2.2 Derivation of continuum model

We now derive a coarse-grained system of partial differential equations describing the evolution of cell density at a larger scale. To do so we take the continuum limit by increasing the number of springs per cell, *m*, while maintaining the total number of cells, *N*, and tissue length, *L*, fixed, and by performing spatial averages over length scales involving a sufficiently large number of cells to define continuous densities, but sufficiently small to retain spatial heterogeneities. We first define the microscopic cell density, 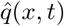, in terms of the spring boundary positions, 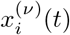, as

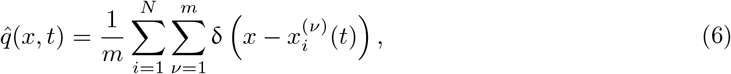

where δ is the Dirac delta function [38,44]. Integrating Equation (6) over the tissue domain, 0 *< x < L*, gives the total number of cells, *N*, which is independent of *m*. We introduce a mesoscopic length scale δ*x* such that 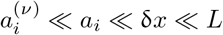 and define a local spatial average which, for the microscopic cell density, 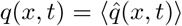, is

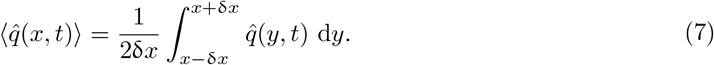

Differentiating Equation (7) with respect to time leads to the general conservation law

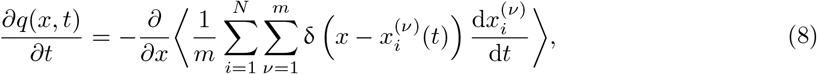

where we use properties of the Dirac delta distribution [44] and take the spatial derivative outside of the average by making use of the fact that δ*x* is small. The averaged term on the right of Equation (8) is the coarse-grained cell density flux, *j*(*x, t*), describing spring migration at the mesoscopic scale, expressed explicitly in terms of the spring boundary positions and velocities [38]. We now introduce three field functions, *f* (*x*, *t*), *k*(*x*, *t*), *a*(*x, t*), for the cell-cell interaction force, the cell stiffness and the resting cell length, respectively, defined such that

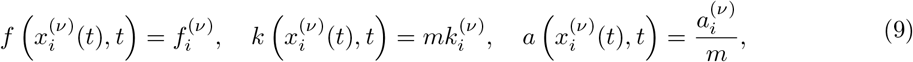

where the scalings for *f*, *k*, and *a*, with respect to *m*, agree with the scalings from the discrete system, see Equation (4). The field functions *k*(*x, t*) and *a*(*x, t*) capture the assumption that spring properties and respective cell properties are constant along spring boundary trajectories, 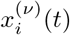. To represent the distribution of spring lengths across the domain, we introduce a continuously differentiable function, *l*(*x, t*), which we define such that

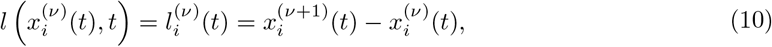

where 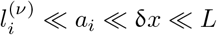. Writing Equation (5) in terms of these continuous variables, expanding each cell-cell interaction force using the small parameter 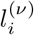, using the viscosity coefficient scaling, and simplifying to leading order gives,

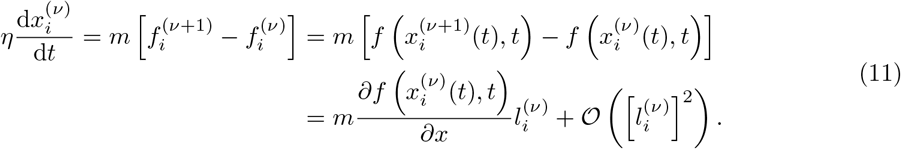

Substituting Equation (11) into Equation (8), relating the spring length to the cell density with 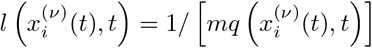, and integrating over the spatial average interval, (*x* − δ*x, x* + δ*x*) gives

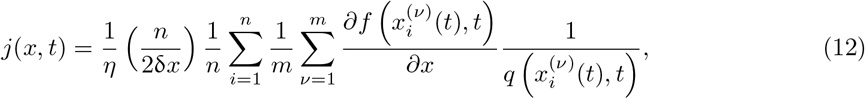

where *n* is the number of cells in the interval (*x −* δ*x, x* + δ*x*) and *i* has been reset to count these cells. Then, taking the limit as *m* → ∞ and performing an average over the *m* springs per cell, gives

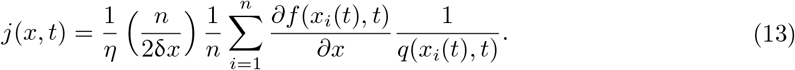

We apply a mean field approximation, as *n* » 1 in (*x* − δ*x, x* + δ*x*) due to *a*_*i*_ « δ*x*, by substituting *q*(*x*_*i*_(*t*), *t*) and *∂f* (*x*_*i*_(*t*), *t*)/*∂x* in the sum with the average density *q*(*x, t*) and the average interaction force gradient *∂f/∂x* in the interval (*x* − δ*x, x* + δ*x*). The factor 1/*q* is now independent of *i* and cancels with the factor *n/*(2δ*x*) which represents the density of cells in the spatial average interval. Then the coarse-grained cell density flux is

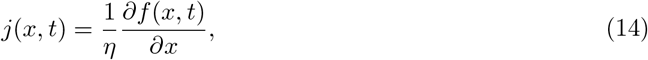

which provides us with an important physical interpretation and is directly related to the velocity, net force and cell-cell interaction force gradient. By inspection of Equation (11) and Equation (14), we see that the cell density flux, *j*, is an advective flux *j* = *qu*, where *u*(*x, t*) = 〈d*x*_*i*_/d*t*〉 is the average velocity induced by the average force gradient 〈*∂f/∂x*〉. We also see that the net force is given by *ηj/q* and the spatially averaged interaction force gradient is given by *ηj*.

Substituting Equation (14) into Equation (8) gives

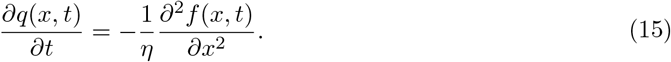

We now return to Equation (9) and differentiate with respect to time to derive an evolution equation for the cell stiffness function

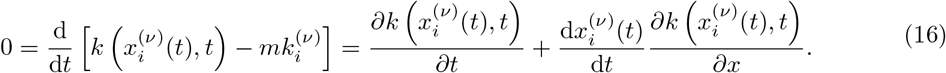

Using Equation (11) and similar developments, the evolution equations for the cell stiffness and resting cell length expressed in terms of mesoscopic variables become

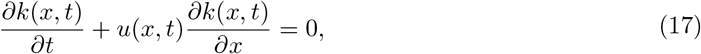

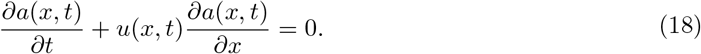

Written in terms of velocity we identify the left-hand sides of Equations (17) and (18) as the convective derivatives of the cell properties.

In summary, the governing equations of the coarse-grained model are given by Equations (15), (17) and (18) with the interaction force *f* given by

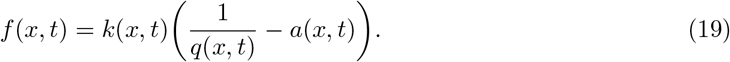

This results in a system of four self-consistent equations for the continuous fields *q*(*x, t*), *k*(*x, t*), *a*(*x, t*), *f* (*x, t*) in terms of spatial position rather than particle trajectories. The initial conditions for the average cell density, cell stiffness and resting cell length are

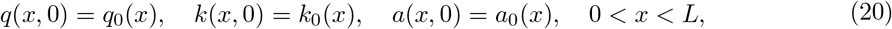

together with no flux boundary conditions for the average cell density, due to the microscopic motion constraints, and Dirichlet boundary conditions for the cell stiffness and resting cell length, as cell properties are constant along cell boundary trajectories,

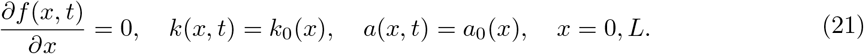

These governing partial differential Equations (15), (17), (18), (19) are solved numerically with the initial conditions (20) and boundary conditions (21), see Supplementary Material Section 2. With homogeneous cell populations the governing equations reduce to the single nonlinear density diffusion equation previously derived in [34],

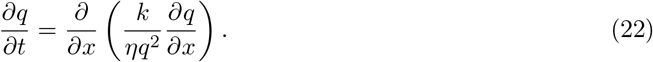

## 3 Results and discussion

In this section we compare solutions of the continuum and discrete models with the expectation that as the number of springs per cell, *m*, increases solutions from the discrete model converge to the corresponding continuum solution.

### 3.1 Homogeneous cell population

We first consider a homogeneous cell population, with one spring per cell, *m* = 1, to illustrate the time evolution of the cell density flux during mechanical relaxation of the tissue. To compare results from the discrete and continuum systems we choose the initial cell configuration (Figure 2(*a*)) to represent a normally distributed cell density,

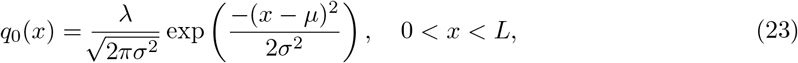

with mean position *µ* = 5 and variance *σ* = 3. We choose *λ* to satisfy 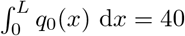 so that with *L* = 10 the total number of cells is *N* = 40, see Supplementary Material Section 1. Then, using the discrete model, we observe that the system relaxes to a uniform cell distribution (Figure 2(*a*)). Figures 2(*b*) and 2(*c*) show how the density and velocity, respectively, propagate along the cell boundary characteristics and demonstrate that the system undergoes temporal relaxation to a steady state configuration. With an eigenmode decomposition of the governing equations of the discrete system, given by Equation (3) and the fixed boundary conditions, we find all eigenvalues are negative which explains the exponential decay behaviour.

**Figure 2:**
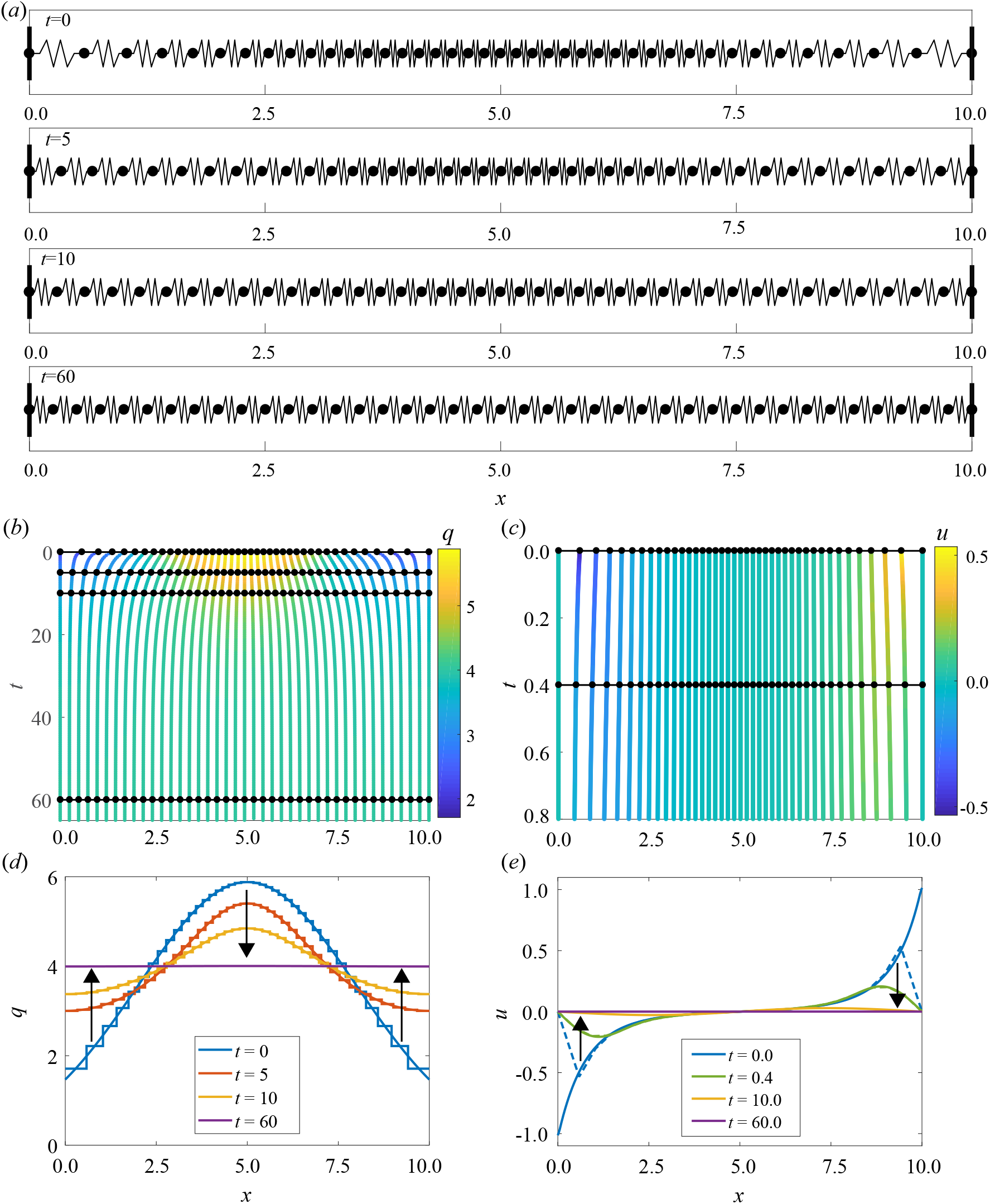
Results for homogeneous *k* and *a*, with *N* = 40 and *m* = 1. (*a*) Snapshots of cell boundary positions and cell lengths at *t* = 0, 5, 15, 60. (*b*) Characteristic diagram for cell boundary position evolution for 0 ≤ *t* ≤ 65. Colour denotes the cell density. Black lines with dots represent snapshots in (*a*) and (*d*). (*c*) Characteristic diagram for cell position evolution for 0.0 ≤ *t* ≤ 0.8. Colour denotes velocity. Black lines and dots represent snapshots in (*e*). (*d*) Cell density snapshots at *t* = 0, 5, 10, 60. Results from discrete/continuum system displayed as stepped/solid lines. (*e*) Velocity snapshots at *t* = 0.0, 0.4, 10.0, 60.0. Results from discrete simulation and continuum system displayed as dashed/solid lines. Arrows indicate the direction of increasing time.

We determine the discrete cell density as the inverse of the spacing between cell boundary trajectories, *q*_*i*_ = 1/(*x*_*i*+1_ − *x*_*i*_) and we assign this value throughout the region *x*_*i*_ < *x* < *x*_*i*+1_. We now compare this discrete information with the density from the continuum system, *q*, obtained by solving Equations (15), (17), (18), and (19). In Figure 2(*d*) we see that the initially normally distributed density tends to the uniform density 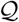, given by 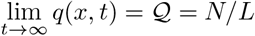, which is independent of *k* and *a*. From Equation (15) we see that this motion is driven by imbalances in the local interaction force field. We relate this to the velocity, *u* = (*∂f/∂x*)/(*ηq*) from Section 2, and we see that as the local imbalances tend to zero the cell boundary velocities tend to zero (Figure 2(*e*)). Due to fast dynamics followed by slow long-term dynamics, results for *t* = 10 and *t* = 60 are mostly overlapping with the steady state (Figure 2(*e*)). This agrees with the interpretation of the discrete system from Equation (3).

### 3.2 Heterogeneous cell population

Here we present results for slowly-varying-in-space and piecewise constant heterogeneous cell populations.

#### 3.2.1 Slowly varying cell population

For slowly-varying-in-space cellular properties, we explore how solutions of the discrete system converge to the solution of the continuum system as *m* increases. We consider heterogeneity in *k* and homogeneous *a* so that, on average, cells are in compression. Figure 3 depicts how the system relaxes to a non-uniform density distribution, due to cell stiffness heterogeneity, as the velocity field *u* tends to zero. From this simulation, we observe higher density in regions of higher *k*. This prediction agrees with the steady state solution to the coarse-grained model, governed by Equations (15), (17), (18), and (19),

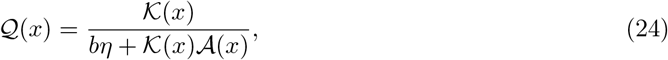

where 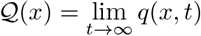, 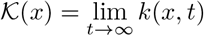 and 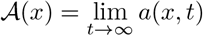, are steady-state solutions and *b* is a constant of integration that is related to *N*. We also observe that, as cell properties are constant along trajectories, the cell stiffness evolves at a fixed location in space. We see in Figure 3(*d-h*) that there is close agreement between the discrete model and the continuum solutions as *m* increases. It is notable that even for low *m* we have excellent agreement between the discrete density and the continuum density at the centre of each spring. However, at spring boundaries the agreement does not hold as well for low *m*. We see similar discrete-continuum agreement when we consider other examples with heterogeneous *k* and homogeneous *a*, with homogeneous *k* and heterogeneous *a*, and heterogeneous *k* and heterogeneous *a* (Supplementary Figures S3-S6).

**Figure 3:**
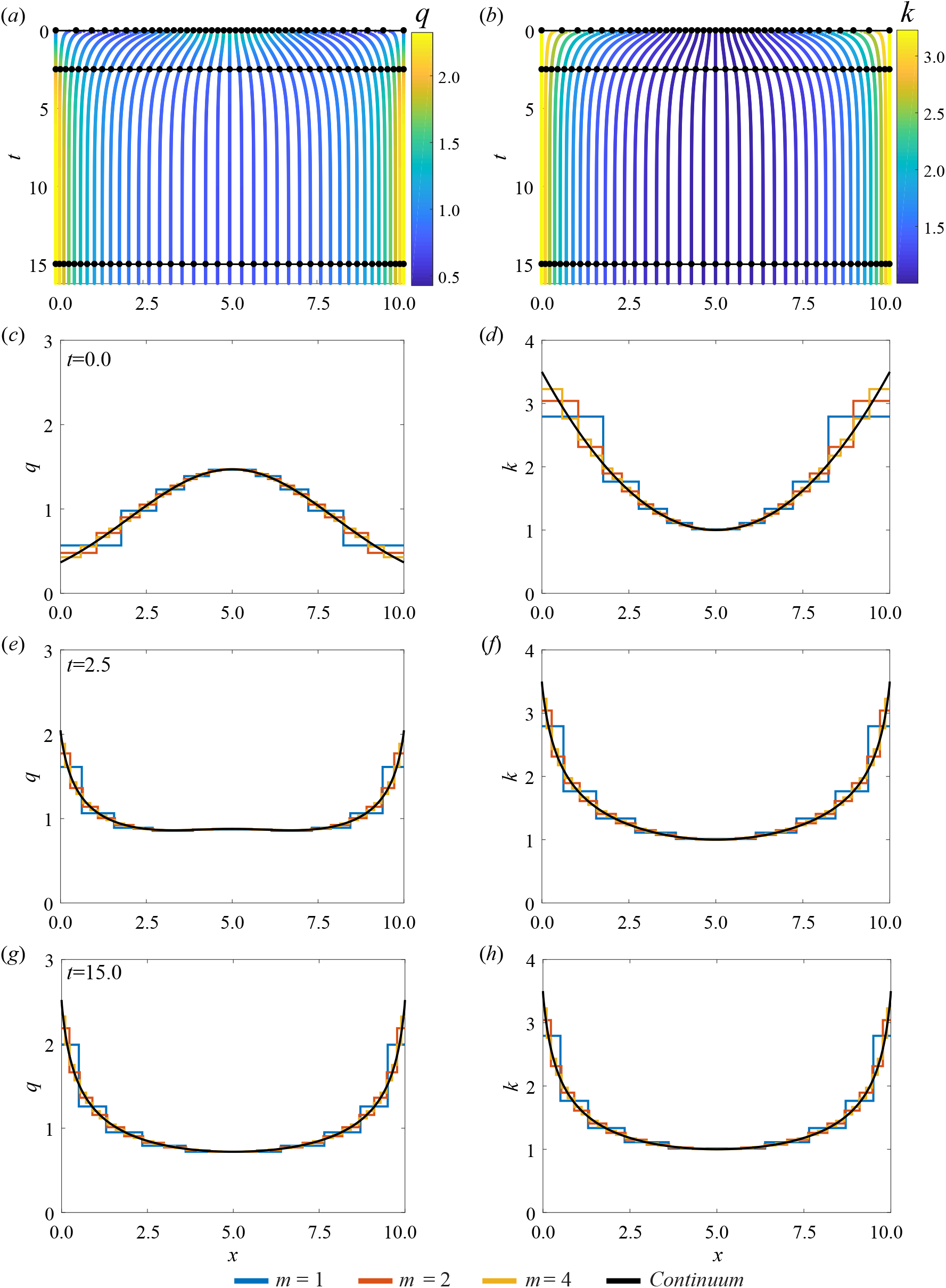
Results for heterogeneous *k* and homogeneous *a* with *N* = 10, *k*_0_(*x*) = 1 + 0.1(*x −* 5)^2^, and *a*_0_(*x*) = 0. (*a,b*) Characteristic diagram for spring boundary position evolution for 0.00 ≤ *t* ≤ 16.25, with *m* = 4 so that every fourth trajectory represents a cell boundary. Colour denotes (*a*) cell density, (*b*) cell stiffness. In (*a,b*) black lines and dots represent times for snapshots in (*c-h*). (*c,e,g*) Cell density snapshots at *t* = 0.0, 2.5, 15.0. (*d,f,h*) Cell stiffness snapshots at *t* = 0.0, 2.5, 15.0. In (*c-h*) lines display results for *N* = 10 with *m* = 1, 2, 4, and continuum system.

#### 3.2.2 Piecewise constant cell population

In this section, we consider a simple scenario with two adjacent tissues, modelled by assuming sharp inhomogeneities in cellular properties. This may represent the boundary between a malignant tissue and a normal tissue. We first explore how solutions from the discrete system converge to the corresponding continuum solution as *m* increases, under these rapidly-varying-in-space conditions. Each tissue has homogeneous cell properties given by cell stiffnesses *k*_1_, *k*_2_ and resting cell lengths *a*_1_, *a*_2_ in the left and right tissue, respectively, with interface position *s*(*t*) (Figure 1(*b*)). For initial conditions, we choose a uniform density, *q*_0_(*x*) = 1, cell properties *k*_1_ = 1/2, *k*_2_ = 1, *a*_1_ = *a*_2_ = 0, *L* = 10 and *s*(0) = 5, respectively. The cell stiffness discontinuity rapidly induces a sharp change in the density at *s*(*t*) followed by slower dynamics until reaching a piecewise constant steady state as *t → ∞* (Figure 4). Even with these sharp inhomogeneities we again observe close agreement between solutions of the discrete and continuum models, especially for the cell stiffness, *k*, where it is difficult to distinguish between the discrete model with different *m* and the solution of the continuum model.

**Figure 4:**
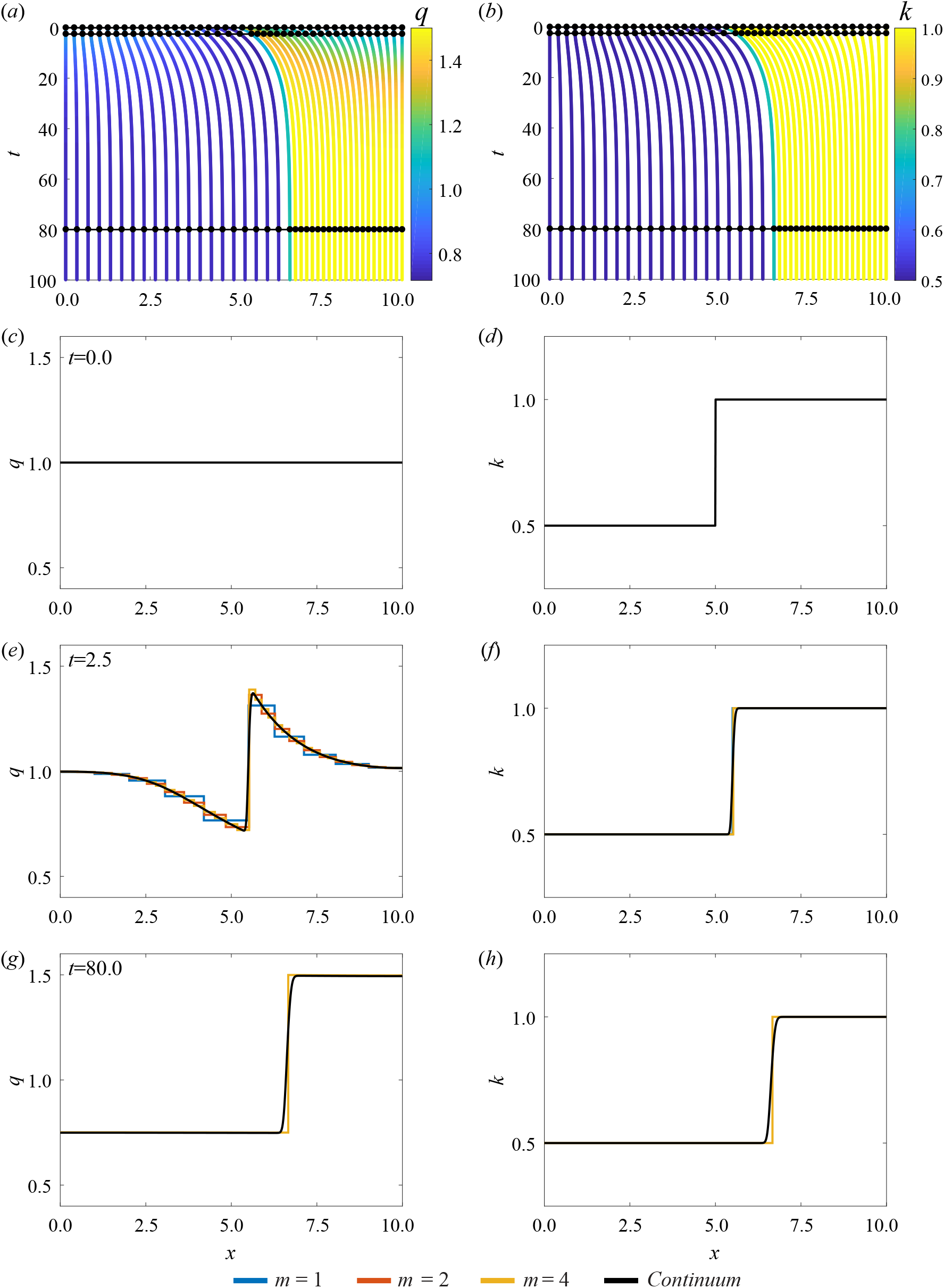
Results for piecewise constant cell properties, with *N* = 10. (*a,b*) Characteristic diagram for spring boundary position evolution for 0 ≤ *t* ≤ 100, with *m* = 4 so that every fourth trajectory represents a cell boundary. Colour denotes (*a*) cell density, (*b*) cell stiffness. In (*a,b*) black lines and dots represent times for snapshots in (*c-h*). (*c,e,g*) Cell density snapshots at *t* = 0.0, 2.5, 80.0. (*d,f,h*) Cell stiffness snapshots at *t* = 0.0, 2.5, 80.0. In (*c-h*) lines display results from *N* = 10 with *m* = 1, 2, 4, and continuum system.

For the cell density, *q*, we again see that agreement at the spring boundaries improves as we increase *m*. This holds especially well given that the numerical discretisation of the continuum model does not explicitly follow the location of the interface, see Supplementary Material Section 2. It could however be determined by evaluating the velocity, d*s*(*t*)/d*t* = *u*, at the interface position.

This simple mechanical relaxation scenario between two tissues enables us to infer some information on the cellular-level properties *k*_*i*_ and *a*_*i*_ by considering the evolution of the interface position, *s*(*t*). The steady state interface position, 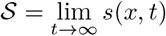 is given by

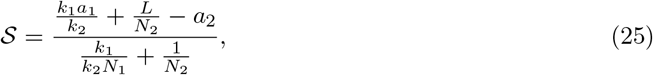

which depends on *k*_1_/*k*_2_, *a*_1_ and *a*_2_. Here *N*_1_ and *N*_2_ represent the total number of cells in the left and right tissues, respectively, see Supplementary Material Section 3. We can identify 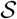 and 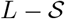 as the lengths of the left and right tissues, respectively, after their mechanical relaxation.

To investigate the influence of *k*_1_/*k*_2_ we vary *k*_1_ and set *k*_2_ = 1. As we have fixed boundaries at *x* = 0 and *x* = *L*, we set *a*_1_ = *a*_2_ = 0 to emphasise properties when we vary *k*_1_, and choose a uniform density initial condition and *N*_1_ = *N*_2_ = 5. Evaluating *s*(*t*) numerically, for efficiency with the discrete model from Equation (3), and 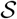 from Equation (25), shows that if *k*_1_ = 0 then 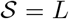 and the left tissue occupies the entire domain. As *k*_1_ → ∞ then 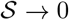 the length of the left tissue decreases (Figures 5(*a,c*)).

**Figure 5:**
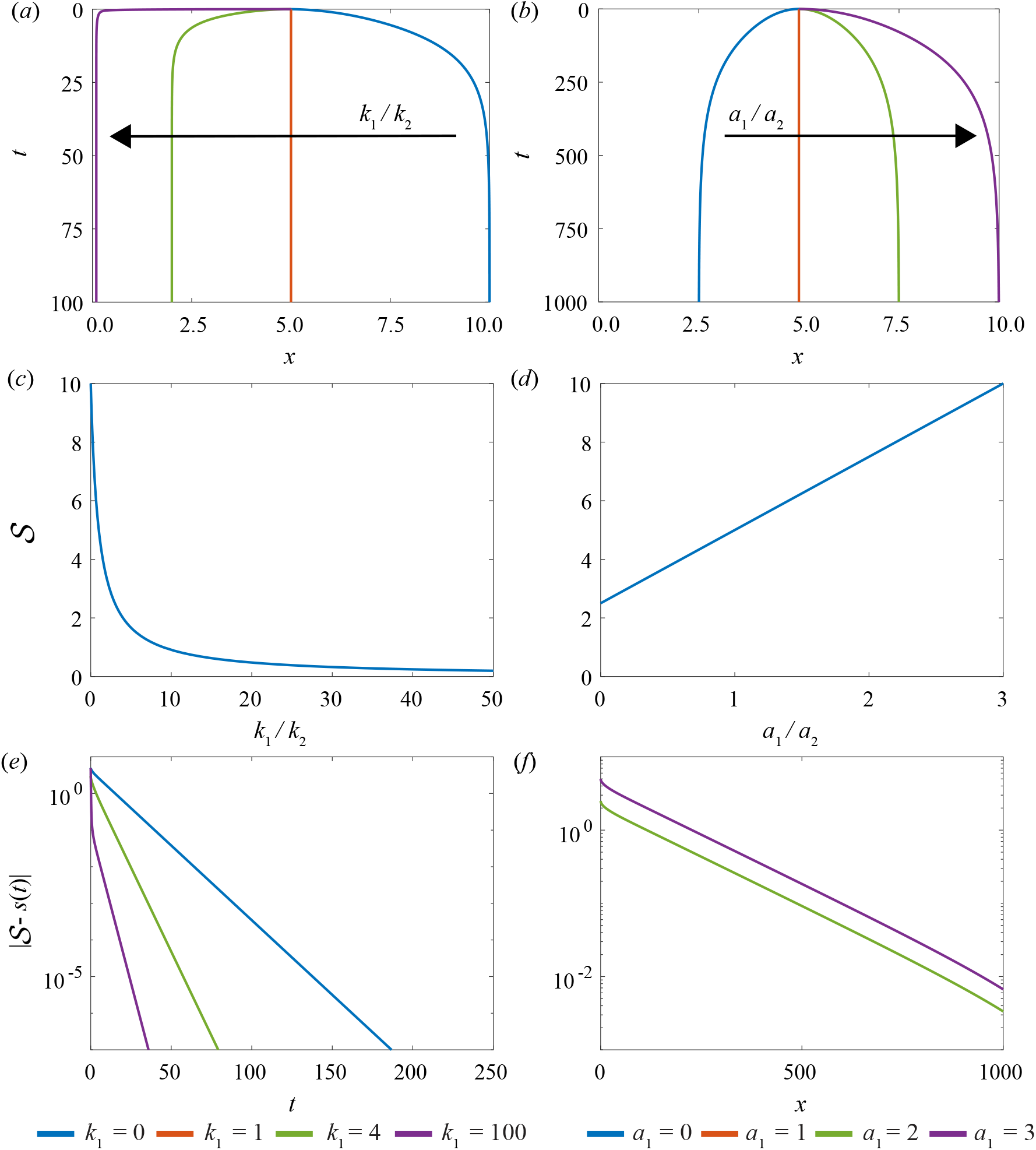
Variation of relative cell stiffness, *k*_1_/*k*_2_, and relative resting cell length, *a*_1_/*a*_2_, in a model with two adjacent tissues and a constant density initial condition. (*a*) Characteristics of the interface position for varying *k*_1_/*k*_2_. The right tissue has fixed cell stiffness *k*_2_ = 1 while the left tissue cell stiffness is varied. (*b*) Characteristics of the interface particle for varying *a*_1_/*a*_2_. The right tissue has fixed resting cell length *a*_2_ = 1 while the resting cell length of the left tissue is varied. Analytical solution for the steady state position of the interface position with given (*c*) relative cell stiffness and (*d*) relative resting cell length. (*e,f*) Absolute difference between position and steady state for interface position for increasing time for varying (*e*) relative cell stiffness and (*f*) relative resting cell length.

Similarly, to investigate the influence of *a*_1_, *a*_2_ we consider *a*_1_/*a*_2_, vary *a*_1_ and set *a*_2_ = 1. We set *k*_1_ = *k*_2_ = 1 which only impacts the rate at which we reach the long-time solution. In contrast to varying *k*_1_/*k*_2_, steady state results depend on the choice of *a*_2_, not just the ratio *a*_1_/*a*_2_, see Equation (25). For example, when *a*_1_ = 0 then 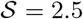 which corresponds to a non-zero minimum left tissue length and a maximum length for the right tissue. We also observe that 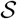 is proportional to *a*_1_ (Figures 5(*b,d*)).

We find that we can use the interface boundary velocities to infer cellular-level properties. Plotting 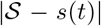 on a logarithmic scale against time shows that we can determine *k*_1_/*k*_2_ from the gradient of the linear section and we can determine *a*_1_/*a*_2_ from the *y*-intercept (Figure 5(*e,f*)). We find that it is easier to distinguish the ratio *k*_1_/*k*_2_ than it is to distinguish the ratio *a*_1_/*a*_2_. If the second tissue was a reference material with known *k*_2_, *a*_2_ we could then determine *k*_1_, *a*_1_.

### 3.3 Case study: breast cancer detection

Recent experiments have proposed a new method to classify breast biopsies in situations where standard histological analysis is inconclusive [6,13,14]. The method is based on determining the stiffness histogram distribution of the tissue using atomic force microscopy. Normal tissues are associated with a single, well-defined unimodal stiffness peak, whereas malignant tissues are associated with a bimodal distribution with a prominent low-stiffness peak. Using our mathematical model, we are able to gain more insight into the differences in mechanical properties of normal and malignant tissues at the cellular level, in particular, the role of the resting cell length, which is not an easy quantity to measure experimentally. It would be impossible to interpret this experimental data with previous models that deal only with homogeneous cell populations.

For this case study, as the experimental data is relatively discrete, we use the discrete model, which we consider to be a sufficiently simple yet insightful portrayal of the biological details. We set the initial state of the system by assuming a uniform initial density distribution and by assigning the cell stiffness of the *i*^th^ cell, *k*_*i*_, so as to reconstruct the unimodal stiffness profile from Figure 1b (top) in [6]. To do so, we normalise the experimental stiffness histogram and interpret the normalised value as the length fraction of the tissue containing stiffness in the given histogram bin (Figure 6(*a*)). This is consistent with the experimental method which implicitly assumes that the probability a cell is examined during a biopsy is proportional to its size [6]. To estimate *k*_*i*_, we randomly sample the unimodal stiffness distribution and arbitrarily assign them to cells *i* = 1, 2, …, *N* in ascending order. Note that the ordering of the cells does not affect our results or the interpretation of our results in any way. We assume *N* = 1000, *m* = 1 and *L* = 10 for illustration purposes. In order for this initial setup to be in equilibrium despite the heterogeneity in stiffness in the tissue, the resting cell lengths *a*_*i*_ must be chosen heterogeneously, per the steady state system of discrete equations, see Supplementary Material Section 4.

**Figure 6:**
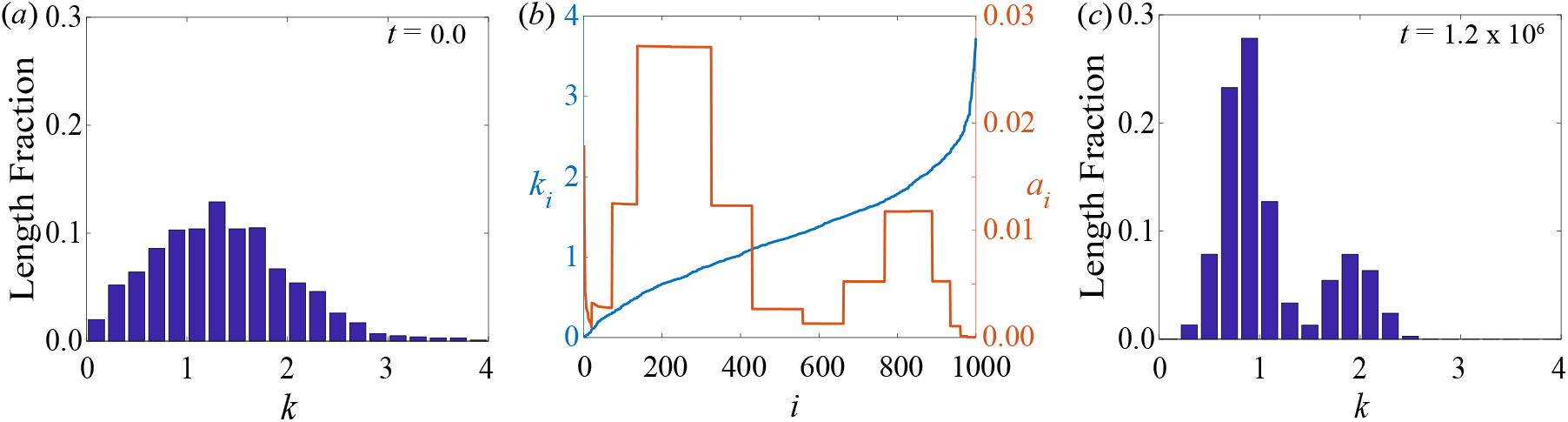
Breast cancer detection case study. (*a*) Initial unimodal stiffness distribution, normalised by tissue length fraction, associated with normal tissues. (*b*) Initial cell stiffness *k*_*i*_ and modified resting cell length *a*_*i*_ for each cell *i* = 1, 2, …,1000, leading to a bimodal stiffness distribution. (*c*) Steady-state stiffness distribution obtained with the modified resting cell lengths, exhibiting a bimodal distribution associated with malignant tissues.

We proceed to consider how a bimodal stiffness distribution, associated with malignant tissues, could arise from such an initial state with a unimodal stiffness distribution. The simplest explanation is that a bimodal stiffness distribution may arise as a result of changes to individual cell stiffnesses, *k*_*i*_, [45] e.g. due to some pre-cancerous biological mechanisms. This model provides an alternative interpretation where the bimodal distribution may arise from changes not solely to the individual cell stiffnesses but to the resting cell lengths also. We now present an extreme case where the bimodal distribution may arise from changes in the resting cell lengths only. Specifically, when we simulate the discrete model with the initial conditions as above, but modify the heterogeneity in the resting cell lengths, *a*_*i*_, to a bimodal profile with high *a*_*i*_ for very low *k*_*i*_, without changing their stiffnesses, *k*_*i*_, the cells redistribute themselves in the tissue in such a way that the tissue stiffness histogram develops a bimodal distribution at mechanical equilibrium (Figure 6(*c*)). We note that this result is not surprising due to the coupling of cell stiffness and resting cell length in the mathematical model. However, this intuitive result may not have been clear had we relied upon experimental data and experimental observations alone. In addition, this approach assumes that cells may have very different lengths which is consistent with biological observations. Specifically, it is understood that breast cancer cells are less stiff and, in general, have a larger diameter in comparison to normal breast cells [46]. This is consistent with other areas of biology, for example, in the context of melanoma biology it is well accepted that cancer cells can be smaller than healthy cells [47,48]. We also note here that changes in the resting cell lengths have been assumed in other works [49] to model two-way feedback between mechanical tensions and signalling and here could similarly represent some unknown underlying pre-cancerous biological mechanisms.

## 4 Conclusion and future work

In this work, we present a one-dimensional cell-based model with heterogeneous cell properties, and its coarse-grained continuum approximation. The motion of cells is driven by cell-cell interaction forces which could represent cell-cell adhesion [41] or compressive stresses [42]. Heterogeneous cell properties, cell stiffness and resting cell length, are constant for each cell and are transported in space by cell movements. The continuum limit is taken by increasing the number of springs per cell, while maintaining the number of cells in the tissue and its fixed total length, and by considering spatial averages over length scales involving a large enough number of cells to define continuous densities but small enough to retain spatial heterogeneities.

Our results shows that solutions of the discrete model approach the solution of the continuum model as the number of springs per cell increases whilst the spring length tends to zero, even for rapidly varying spatial cell properties. Excellent agreement is observed even for few springs per cell at the centre of each cell. For the examples presented in this work, we find that the solution of the discrete model can be obtained much faster than the solution of the continuum model. However, the time required to simulate the discrete model increases rapidly with the number of cells. In contrast, the time required to simulate the continuum model is independent of the number of cells. Therefore, when we have large numbers of cells, as in an epithelial tissue, the continuum model is advantageous. Another advantage of the continuum model is that we can quickly develop exact closed form expressions for the long-time interface position which are more difficult to establish with the discrete model. Furthermore, the continuum model allows us to understand macroscale phenomena which are not obvious from microscopic interactions. The fact that the cell density flux in the continuum model, a macroscopic quantity, is explicitly related to the gradient of the cell-cell interaction force may have been anticipated, but it is not obvious from the microscopic interactions that this leads to an effective non-linear diffusive transport. Finally, because the continuum model exhibits explicit relationships between macroscopic quantities, it will be more useful for inverse problems.

By dealing explicitly with heterogeneous cell populations, this model has many potential applications. The first application we consider is a simple tissue relaxation simulation, where we track the position of the interface between two distinct adjacent tissues as the system mechanically relaxes, to infer cellular-level properties. Results suggest it is easier to determine the relative cell stiffnesses than it is to determine the relative resting cell lengths. Results also show that when cells are, on average, in tension a tissue with lower stiffness extends and compresses a tissue with higher stiffness. In the second application, we use the model to interpret recent experiments in breast cancer detection which reveal distinct stiffness profiles associated with normal, benign and malignant tissues [6]. We show that a bimodal stiffness distribution, associated with a malignant tissue, could arise from a unimodal stiffness distribution, associated with a normal tissue, from changes not just in cell stiffnesses but from changes in the resting cell length’s only. The resting cell length is not an easily measured experimental quantity and these results suggest that this could be an important variable to consider.

Many extensions of this work are possible, both mathematically and biologically. Important extensions will be to introduce cell proliferation, apoptosis, and free boundaries where the continuum limit is less obvious [34,39,40]. Another interesting extension will be to generalise the cell-cell interaction force law to include nonlinear effects for large separations [35,40]. These extensions will be the subject of future works. Finally, the model’s ability to relate cellular-level stiffness data and tissue-level information has many potential extensions biologically including applying the model to particular scenarios such as epithelial tissue mechanics, cancer progression [4,5], cancer detection [6–8], wound healing [10], and morphogenesis [11].

### Data Access

This article does not contain any additional data. Key algorithms used to generate results are available on GitHub.

## Author Contributions

All authors conceived and designed the study; R.J.M. performed numerical simulations and drafted the article; all authors provided comments and gave final approval for publication.

## References

[1] Trepat X, Sahai E. 2018 Mesoscale physical principles of collective cell organisation. Nat. Phys. 14. 671–682. (doi:10.1038/s41567-018-0194-9)

[2] Altschuler SJ, Wu LF. 2010 Cellular heterogeneity: when do differences make a difference? Cell. 141. 559–563. (doi:10.1016/j.cell.2010.04.033)

[3] Gallaher JA, Enriquez-Navas PM, Luddy KA, Gatenby RA, Anderson ARA. 2018 Spatial heterogeneity and evolutionary dynamics modulate time to recurrence in continuous and adaptive cancer therapies. Cancer Res. 78. 2127–2139. (doi:10.1158/0008-5472.CAN-17-2649)

[4] Samuel MS, Lopez JI, McGhee EJ, Croft DR, Strachan D, Timpson P, Munro J, Schröder E, Zhou J, Brunton, VJ, et al. 2011 Actomyosin-mediated cellular tension drives increased tissue stiffness and β-catenin activation to induce epidermal hyperplasia and tumor growth. Cancer Cell. 19. 776–791. (doi:10.1016/j.ccr.2011.05.008)

[5] Nguyen AV, Nyberg KD, Scott MB, Welsh AM, Nguyen AH, Wu N, Hohlbauch SV, Geisse NA, Gibb EA, Robertson AG, et al. Stiffness of pancreatic cancer cells is associated with increased invasive potential. Integr. Biol. 8. 1232–1245. (doi:10.1039/c6ib00135a)

[6] Plodinec M, Loparic M, Monnier CA, Obermann EC, Zanetti-Dallenbach R, Oertle P, Hyotyla JT, Aebi U, Bentires-Alj M, Lim RYH, et al. 2012 The nanomechanical signature of breast cancer. Nat. Nanotechnol. 7. 757–764. (doi:10.1038/nnano.2012.167)

[7] Huo CW, Chew GL, Britt KL, Ingman WV, Henderson MA, Hopper JL, Thompson EW. 2014 Mammographic density-a review on the current understanding of its association with breast cancer. Breast Cancer Res. Tr. 144. 479–502. (doi:10.1007/s10549-014-2901-)

[8] Suresh S. 2007 Elastic clues in cancer detection. Nat. Nanotechnol. 2. 748–749. (doi:10.1038/nnano.2007.397)

[9] Alibert C, Goud, B, Manneville J-B. 2017 Are cancer cells really softer than normal cells? Biol. Cell. 109. 167–189. (doi:10.1111/boc.201600078)

[10] Evans ND, Oreffo ROC, Healy E, Thurner PJ, Man YH. 2013 Epithelial mechanobiology, skin wound healing, and the stem cell niche. J. Mech. Behav. Biomed. 28. 397–409. (doi:10.1016/j.jmbbm.2013.04.023)

[11] Fletcher AG, Cooper F, Baker RE. 2017 Mechanocellular models of epithelial morphogenesis. Phil. Trans. R. Soc. B. 372. 20150519. (doi:10.1098/rstb.2015.0519)

[12] Levental I, Georges PC, Janmey PA. 2006 Soft biological materials and their impact on cell function. Soft Matter. 3. 299–306. (doi:10.1039/b610522j)

[13] Lekka M, Gil D, Pogoda K, Duliǹska-Litewka J, Jach R, Gostek J, Klymenko O, Prauzner-Bechcicki S, Stachura Z, Wiltowska-Zuber J, et al. 2012 Cancer cell detection in tissue sections using AFM. Arch. Biochem. Biophys. 518. 151–156. (doi:10.1016/j.abb.2011.12.013)

[14] Lekka M. 2012 Atomic force microscopy: a tip for diagnosing cancer. Nat. Nanotechnol. 7. 691–692. (doi:10.1038/nnano.2012.196)

[15] Lekka M. 2016 Discrimination between normal and cancerous cells using AFM. BioNanoSci. 6. 65–80. (doi:10.1007/s12668-016-0191-3)

[16] Cross SE, Jin Y, Rao J, Gimzewski JK. 2007 Nanomechanical analysis of cells from cancer patients. Nat. Nanotechnol. 2. 780–783. (doi:10.1038/nnano.2007.388)

[17] Pathmanathan P, Cooper J, Fletcher A, Mirams G, Murray P, Osborne J, Pitt-Francis J, Walter A, Chapman SJ. 2009 A computational study of discrete mechanical tissue models. Phys. Biol. 6. 036001. (doi:10.1088/1478-3975/6/3/036001)

[18] Osborne JM, Fletcher AG, Pitt-Francis JM, Maini PK, Gavaghan DJ. 2017 Comparing individual-based approaches to modelling the self-organization of multicellular tissues. PLoS Comput. Biol. 13. e1005387. (doi:10.1371/journal.pcbi.1005387)

[19] Sandersius SA, Weijer CJ, Newman TJ. 2011 Emergent cell and tissue dynamics from sub-cellular modeling of active biomechanical processes. Phys. Biol. 8. 045007. (doi:10.1088/1478-3975/8/4/045007)

[20] Ingber DE. 2003 Tensegrity I. Cell structure and hierarchical systems biology. J. Cell Sci. 116. 1157–1173. (doi:10.1242/jcs.00359)

[21] Armstrong NJ, Painter KJ, Sherratt JA. 2006 A continuum approach to modelling cell-cell adhesion. J. Theor. Biol. 243. 98–113. (doi:10.1016/j.jtbi.2006.05.030)

[22] Collis J, Brown DL, Hubbard ME, O’Dea RD. 2017 Effective equations governing an active poroelastic medium. Proc. R. Soc. A. 473. 20160755. (doi:10.1098/rspa.2016.0755)

[23] Penta R, Ambrosi D, Shipley RJ. 2013 Effective governing equations for poroelastic growing media. Q. J. Mech. Appl. Math. 67. 69–91. (doi:10.1093/qjmam/hbt024)

[24] Osborne JM, Walter A, Kershaw SK, Mirams GR, Fletcher AG, Pathmanathan P, Gavaghan D, Jensen OE, Maini PK, Byrne HM. 2010 A hybrid approach to multi-scale modelling of cancer. Phil. Trans. R. Soc. A. 368. 5013–5028. (doi:10.1098/rsta.2010.0173)

[25] Anderson ARA. 2005 A hybrid mathematical model of solid tumour invasion: the importance of cell adhesion. Math. Med. Biol. 22. 163–186. (doi:10.1093/imammb/dqi005)

[26] Vavourakis V, Wijeratne PA, Shipley R, Loizidou M, Stylianopoulos T, Hawkes DJ. 2017 A validated multiscale in-silico model for mechano-sensitive tumour angiogenesis and growth. PLoS Comput. Biol. 13. e1005259. (doi:10.1371/journal.pcbi.1005259)

[27] Rajagopal V, Holmes WR, Lee PVS. 2018 Computational modeling of single-cell mechanics an cytoskeletal mechanobiology. Wires. Syst. Biol. Med. 10. 5–7. (doi:10.1002/wsbm.1407)

[28] Tambe DT, Hardin CC, Angelini TE, Rajendran K, Park CY, Serra-Picamal X, Zhou EH, Zaman MH, Butler JP, Weitz DA, et al. 2011 Collective cell guidance by cooperative intercellular forces. Nat. Mater. 10. 469–475. (doi:10.1038/nmat3025)

[29] Fozard JA, Byrne HM, Jensen OE, King JR. 2010 Continuum approximations of individual-based models for epithelial monolayers. Math. Med. Biol. 27. 39–74. (doi:10.1093/imammb/dqp015)

[30] O’Dea RD, King JR. 2012 Continuum limits of pattern formation in hexagonal-cell monolayers. J. Math. Biol. 64. 579–610. (doi:10.1007/s00285-011-0427-3)

[31] Middleton AM, Fleck C, Grima R. 2014 A continuum approximation to an off-lattice individual-cell based model of cell migration and adhesion. J. Theor. Biol. 359. 220–232. (doi:10.1016/j.jtbi.2014.06.011)

[32] Matsiaka OM, Penington CJ, Baker RE, Simpson MJ. 2018 Discrete and continuum approximations for collective cell migration in a scratch assay with cell size dynamics. B. Math. Biol. 80. 738–757. (doi:10.1007/s11538-018-0398-2)

[33] Bodnar M, Velazquez JJL. 2005 Derivation of macroscopic equations for individual cell-based models: a formal approach. Math. Method Appl. Sci. 28. 1757–1779. (doi:10.1002/mma.638)

[34] Murray PJ, Edwards CM, Tindall MJ, Maini PK. 2009 From a discrete to a continuum model of cell dynamics in one dimension. Phys. Rev. E. 80. 031912. (doi:10.1103/PhysRevE.80.031912)

[35] Murray PJ, Edwards CM, Tindall MJ, Maini PK. 2012 Classifying general nonlinear force laws in cell-based models via the continuum limit. Phys. Rev. E. 85. 021921. (doi:10.1103/PhysRevE.85.021921)

[36] Murray PJ, Kang J, Mirams GR, Shin S-Y, Byrne HM, Maini PK, Cho KH. 2010 Modelling spatially regulated β-catenin dynamics and invasion in intestinal crypts. Biophys. J. 99. 716–725. (doi:10.1016/j.bpj.2010.05.016)

[37] Murray PJ, Walter A, Fletcher AG, Edwards CM, Tindall MJ, Maini PK. 2011 Comparing a discrete and continuum model of the intestinal crypt. Phys. Biol. 8. 026011. (doi:10.1088/1478-3975/8/2/026011)

[38] Evans DJ, Morriss, G. 2008 Statistical Mechanics of Nonequilibrium Liquids. Cambridge, UK: Cambridge University Press.

[39] Lorenzi T, Murray PJ, Ptashnyk M. 2019 From individual-based mechanical models of multi-cellular systems to free-boundary problems. arXiv Preprint. (arXiv:10.1016/1903.06590)

[40] Baker RE, Parker A, Simpson MJ. 2018 A free boundary model of epithelial dynamics. J. Theor. Biol. (doi:10.1016/j.jtbi.2018.12.025)

[41] Johnston ST, Simpson MJ, Plank MJ. 2013 Lattice-free descriptions of collective motion with crowding and adhesion. Phys. Rev E. 88. 062720. (doi:10.1103/PhysRevE.88.062720)

[42] Tse JM, Cheng G, Tyrrell JA, Wilcox-Adelman SA, Boucher Y, Jain RK, Minn LL. 2011 Mechanical compression drives cancer cells toward invasive phenotype. P. Natl. Acad. Sci. USA. 109. 911–916. (doi:10.1073/pnas.1118910109)

[43] Meineke FA, Potten CS, Loeffler M. 2001 Cell migration and organisation in the intestinal crypt using a lattice-free model. Cell Prolif. 34. 253–266. (doi:10.1046/j.0960-7722.2001.00216.x)

[44] Lighthill MJ. 1958. An Introduction to Fourier Analysis and Generalised Functions. Cambridge, UK: Cambridge University Press.

[45] Guck J, Schinkinger S, Lincoln B, Wottawah F, Ebert S, Romeyke M, Lenz D, Erickson HM, Ananthakrishnan R et al. 2005 Optical deformability as an inherent cell marker for testing malignant transformation and metastatic competence. Biophys. J. 88. 6893698. (doi:10.1529/biophysj.104.045476)

[46] TruongVo TN, Kennedy RM, Chen H, Chen A, Berndt A, Agarwal M, Zhu L, Nakshatri H, Wallace J, Na S et al. 2017 Microfluidic channel for characterizing normal and breast cancer cells. J. Micromech. Microeng. 27. 035017. (doi:10.1088/1361-6439/aa5bbb)

[47] Haridas P, Penington CJ, McGovern JA, McElwain DLS. 2017 Quantifying rates of cell migration and cell proliferation in co-culture barrier assays reveals how skin and melanoma cells interact during melanoma spreading and invasion. J. Theor. Biol. 423. 13–25. (doi:10.1016/j.jtbi.2017.04.017)

[48] Haridas P, McGovern JA, McElwain DLS. 2017 Quantitative comparison of the spreading and invasion of radial growth phase and metastatic melanoma cells in an three-dimensional human skin equivalent model. PeerJ 5. e3754. (doi:10.7717/peerj.3754)

[49] Zmurchok C, Bhaskar D, Edelstein-Keshet L. 2018 Coupling mechanical tension and GTPase signaling to generate cell and tissue dynamics. Phys. Biol. 15. 046004. (doi:10.1088/1478-3975/aab1c0)

