## Supplementary Information for "A one-dimensional individual-based mechanical model of cell movement in heterogeneous tissues and its coarse-grained approximation"

**S5 Supplementary Figures** **7**

### S1 Discrete model simulation

In this section, we describe how to perform simulations with the discrete model.

#### S1.1 Discrete equations

Here, we describe the method to solve Equation (1), governing the discrete model, with $m$ springs per cell, for the position of each spring boundary, $x_i^{(\nu)}$, for $i = 1, 2, \ldots, N$ and $\nu = 1, 2, \ldots, m$ and $x_{N+1}^{(1)}$. The equations are included here for convenience:

$$\eta^{(\nu)} \frac{\mathrm{d}x_i^{(\nu)}}{\mathrm{d}t} = f_i^{(\nu+1)} - f_i^{(\nu)}; \tag{1}$$

$$f_i^{(\nu)} = k_i^{(\nu-1)} \left( x_i^{(\nu)} - x_i^{(\nu-1)} - a_i^{(\nu-1)} \right). \tag{2}$$

#### S1.2 Converting cell density into initial positions

We now explain how to convert an initial distribution of density, $q_0(x)$, into an initial condition for the discrete model by determining the initial spring boundary positions, $x_i^{(\nu)} = x_i^{(\nu)}(0)$, for $i = 1, 2, \ldots, N$ and $\nu = 1, 2, \ldots, m$, and $x_{N+1}^{(1)}$. To solve for these positions we use the Matlab fsolve function [1] applied to the system of the equations

$$x_1^{(1)} = 0, \tag{3}$$

$$\frac{\dfrac{1}{x_i^{(\nu-1)} - x_i^{(\nu)}} + \dfrac{1}{x_i^{(\nu)} - x_i^{(\nu+1)}}}{\dfrac{x_i^{(\nu+1)} - x_i^{(\nu-1)}}{2}} - m \left. \frac{\partial q_0(x)}{\partial x} \right|_{x = x_i^{\nu}} = 0, \tag{4}$$

$$x_{N+1}^{(1)} = L. \tag{5}$$

Equations (3) and (5) arise from the fixed boundary conditions. Equation (4) arises from equating the approximate numerical gradient of the density from the discrete system at a position $x_i^{(\nu)}$ with the gradient of $q_0(x)$ at the same position. To evaluate the numerical gradient we use the midpoints of the domains $x_i^{(\nu-1)} < x < x_i^{(\nu)}$ and $x_i^{(\nu)} < x < x_{i+1}^{(\nu+1)}$. The density in the first domain is given by $1/\left[m\left(x_i^{(\nu)} - x_i^{(\nu-1)}\right)\right]$ and similarly $1/\left[\left(m(x_i^{(\nu+1)} - x_i^{(\nu)}\right)\right]$ for the second domain (Figure S1).

#### S1.3 Assigning spring properties

In this section we explain our approach to assigning spring properties assuming that we know the initial cell boundary positions, $x_i^{(\nu)}$, the initial cell stiffness distribution, $k_0(x)$, and initial resting cell length distribution, $a_0(x)$. Here we consider the cell stiffness initial condition and note that the same ideas apply to assign the resting cell length. We consider spring $\nu$ in cell $i$, located at $x_i^{(\nu)} < x < x_i^{(\nu+1)}$; let $\widetilde{x}_i^{(\nu)}$ be the position of the median of $k_0(x)$ in this domain, and use this to define the discrete spring stiffness as $k_i^{(\nu)} = mk_0\left(\widetilde{x}_i^{(\nu)}\right)$ (Figure S2). For the resting cell length this

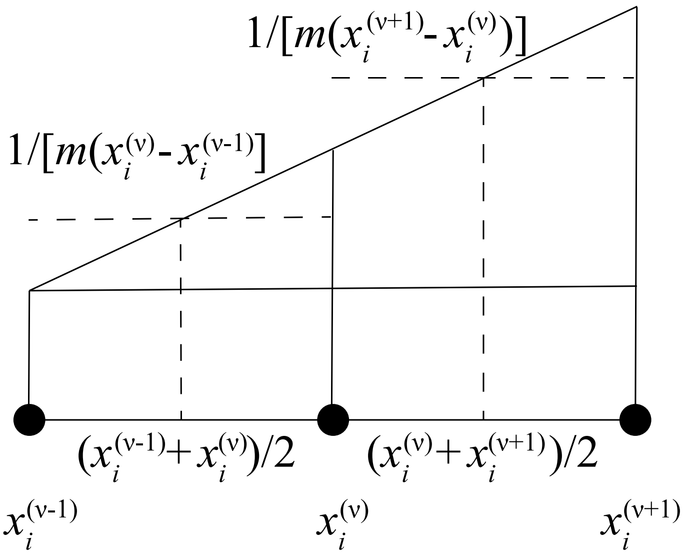

Figure S1: Schematic to determine the approximate numerical gradient of the density from the discrete system at position $x_i^{\nu}$.

would be $a_i^{(\nu)} = a_0\left(\widetilde{x}_i^{(\nu)}\right)/m$. This median position, $\widetilde{x}_i^{(\nu)}$, is solved for by equating the integral of the initial condition spring stiffness function in $x_i^{(\nu)} < x < \widetilde{x}_i^{(\nu)}$ and $\widetilde{x}_i^{(\nu)} < x < x_i^{(\nu+1)}$,

$$\int_{x_i^{(\nu)}}^{\widetilde{x}_i^{(\nu)}} k_0(x)\ \mathrm{d}x = \int_{\widetilde{x}_i^{(\nu)}}^{x_i^{(\nu+1)}} k_0(x)\ \mathrm{d}x. \tag{6}$$

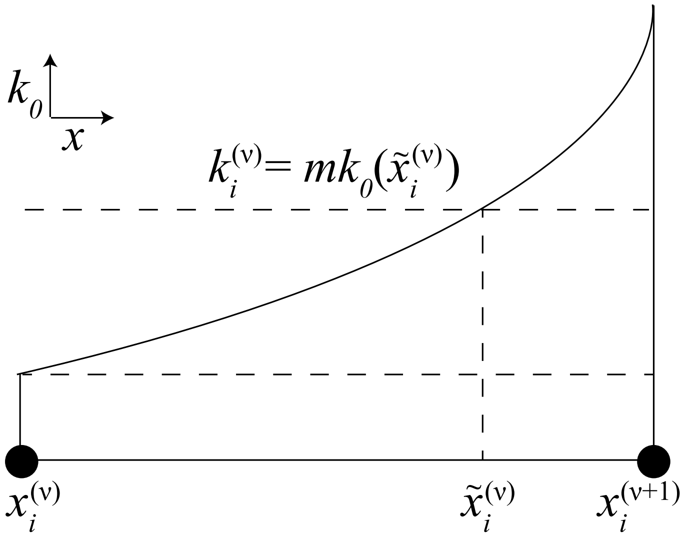

Figure S2: Schematic to determine the position $\widetilde{x}_i^{(\nu)}$ used to define the discrete spring stiffness value for spring $\nu$ in cell $i$, $k_i^{(\nu)} = mk_0\left(\widetilde{x}_i^{(\nu)}\right)$.

##### S1.4 Numerical methods

This discrete model, with $m$ springs per cell, is governed by Equations (1) with the fixed boundary conditions $x_1^{(1)} = 0$, $x_{N+1}^{(1)} = L$. Appropriate scalings of the cell properties are required to determine the spring properties, $k_i^{(\nu)} = mk_0\left(\tilde{x}_i^{(\nu)}\right)$ and $a_i^{(\nu)} = a_0\left(\tilde{x}_i^{(\nu)}\right)/m$. The viscosity coefficient must also be scaled appropriately through $\eta^{(\nu)} = \eta/m$. These form a system of $Nm - 1$ ordinary differential equations, with the two boundary conditions, and initial conditions for the positions, $x_i^{(\nu)}$, spring stiffnesses, $k_i^{(\nu)}$, and resting spring lengths, $a_i^{(\nu)}$, of each spring $\nu$ in cell $i$, and viscosity coefficient for the system, $\eta^{(\nu)}$. We solve this system using Matlab ode15s [2]. Alternatively, this system can be analysed with an eigenmode decomposition which is convenient to determine the steady state.

#### S2 Continuum model simulation

Here we describe the numerical method used to solve the partial differential equations associated with the continuum description.

##### S2.1 Discretisation scheme

For convenience we re-state the governing equations:

$$\frac{\partial q(x,t)}{\partial t} = -\frac{1}{\eta}\frac{\partial^2 f(x,t)}{\partial x^2}; \tag{7}$$

$$\frac{\partial k(x,t)}{\partial t} = -\frac{1}{\eta}\frac{1}{q(x,t)}\frac{\partial f}{\partial x}\frac{\partial k(x,t)}{\partial x}; \tag{8}$$

$$\frac{\partial a(x,t)}{\partial t} = -\frac{1}{\eta}\frac{1}{q(x,t)}\frac{\partial f}{\partial x}\frac{\partial a(x,t)}{\partial x}; \tag{9}$$

$$f(x,t) = k(x,t)\left(\frac{1}{q(x,t)} - a(x,t)\right). \tag{10}$$

The first step is to substitute the interaction force from Equation (10) into Equations (7), (8) and (9). We solve Equation (7) implicitly, and we solve Equations (8) and (9) for the cell stiffness and resting cell length, respectively, explicitly [4]. First, we uniformly discretise the domain with nodes spaced $\Delta x$ apart. The nodes are indexed $j = 1, \ldots, R$, where $R$ is the total number of spatial nodes. We apply an upwinding scheme using a numerically determined velocity at each node, which is defined for node $j$ and time step $n$ as $v_j^n$, given by

$$v_j^n = \frac{1}{\eta}\frac{1}{q_j^n}\left[k_{j+1}^n\left(\frac{1}{q_{j+1}^n} - a_{j+1}^n\right) - k_{j-1}^n\left(\frac{1}{q_{j-1}^n} - a_{j-1}^n\right)\right]. \tag{11}$$

We use non-constant time stepping for efficiency with the timestep to advance the numerical solution from timestep $n$ to timestep $n+1$ denoted $\Delta t^n$. The value of this timestep is determined based on the maximum numerical velocity across all nodes at time $n$, $\max_j(v_j^n)$, and is chosen as $\Delta t^n = \min\left(0.001(\Delta x)^2, 0.00001(\Delta x)^2/\max_j(v_j^n)\right)$, where $\Delta x = 0.01$ to produce Figure 4 and

Supplementary Figures S3, S4 and S6, and $\Delta x = 0.05$ otherwise.

We solve Equation (7) using a Crank-Nicolson approximation. At the central nodes we have

$$\begin{aligned}\frac{q_j^{n+1}-q_j^n}{\Delta t^n} =& -\frac{1}{2}\frac{1}{(\Delta x)^2}\left[\frac{k_{j+1}^n}{\eta}\left(\frac{1}{q_{j+1}^n}-a_{j+1}\right)-2\frac{k_j^n}{\eta}\left(\frac{1}{q_j^n}-a_j\right)+\frac{k_{j-1}^n}{\eta}\left(\frac{1}{q_{j-1}^n}-a_{j-1}\right)\right]\\ &-\frac{1}{2}\frac{1}{(\Delta x)^2}\left[\frac{k_{j+1}^{n+1}}{\eta}\left(\frac{1}{q_{j+1}^{n+1}}-a_{j+1}\right)-2\frac{k_j^{n+1}}{\eta}\left(\frac{1}{q_j^{n+1}}-a_j\right)+\frac{k_{j-1}^{n+1}}{\eta}\left(\frac{1}{q_{j-1}^{n+1}}-a_{j-1}\right)\right],\\ & j=3,\ldots,R-2,\ n=0,\ldots T,\end{aligned} \tag{12}$$

where we use the approximation

$$\frac{1}{q_j^{n+1}} = \frac{q_j^{n+1}}{\left(q_j^n\right)^2}, \tag{13}$$

for the terms at timestep $n+1$ on the right-hand side of Equation (12). This approximation allows us to write the discretised system of equations in tridiagonal form. For the boundary condition at $x = 0$, corresponding to node $j = 1$, we apply a second order forward difference stencil so that

$$q_1^n = 1\Big/\left[a_1^n + \frac{4k_2^n}{3k_1^n}\left(\frac{1}{q_2^n}-a_2^n\right)-\frac{k_3^n}{3k_1^n}\left(\frac{1}{q_3^n}-a_3^n\right)\right]. \tag{14}$$

To obtain the equation for node $j = 2$ we set $j = 2$ in Equation (12) and replace $q_1^n$ with Equation (14). Similarly, to obtain an equation for node $j = R$ at the right boundary $x = L$ we apply a second order backwards difference stencil,

$$q_R^n = 1\Big/\left[a_R^n - \frac{k_{R-2}^n}{3k_R^n}\left(\frac{1}{q_{R-2}^n}-a_{R-2}^n\right)+\frac{4k_{R-1}^n}{3k_R^n}\left(\frac{1}{q_{R-1}^n}-a_{R-1}^n\right)\right]. \tag{15}$$

This allows us to form an equation for node $j = R-1$ also. Now we use the Thomas algorithm [3] to advance one time step for the equations governing nodes $j = 2,\ldots,R-1$. Using the results for nodes $j = 2,\ldots,R-1$ and Equations (14) and (15) we can update the boundary nodes $q_1^{n+1}, q_R^{n+1}$.

While performing each temporal step we must also update the cell stiffnesses, $k_j^n$, and resting cell lengths, $a_j^n$, at each node. At the boundaries, $x = 0$ and $x = L$, corresponding to nodes $j = 1$ and $j = R$, these cell properties are fixed so $k_1^n = k_1^0$ and $a_1^n = a_1^0$ for $n = 1,\ldots,T$. For the interior nodes $j = 2,\ldots,R-1$, we apply an explicit method with upwinding. The sign of $\max_j(v_j^n)$ determines whether we apply forward or backward difference stencils. For example, if $\max_j(v_j^n) > 0$ then we apply a backward first order difference to the cell stiffness, Equation (8),

$$\frac{k_j^{n+1}-k_j^n}{\Delta t^n} = -\frac{1}{\eta}\frac{k_j^n-k_{j-1}^n}{(\Delta x)^2}\left[k_j^n\left(\frac{1}{q_j^n}-a_j^n\right)-k_{j-1}^n\left(\frac{1}{q_{j-1}^n}-a_{j-1}^n\right)\right],\ j=2,\ldots,R-1. \tag{16}$$

Cell properties are constant along cell boundary trajectories. Therefore, if we have a cell property which is initially homogeneous it will remain homogeneous. The numerical method can then be simplified by not simulating the related cell property equation and replacing its appearance in the other equations with its constant value.

### S3 Steady state analysis for two tissue model

We obtain an analytical expression for the steady state position of the interface, $\mathcal{S}$, between two distinct adjacent tissues. Suppose that the left-most tissue is characterised by $N_1$ cells with stiffness $k_1$ and resting spring length $a_1$, for $0 < x < \mathcal{S}$. Similarly, suppose that the right-most tissue is characterised by $N_2$ cells with stiffness $k_2$ and resting spring length $a_2$ for $\mathcal{S} < x < L$. Considering the continuum system given by Equations (7), (8), (9), and (10) for each tissue at steady state, the equations governing the evolution of the cellular properties are trivially solved as we have a homogeneous cell population in each tissue. However, solving Equation (7) and applying the no flux density boundary conditions gives, for the first tissue,

$$\frac{k_1}{\eta}\left(\frac{1}{q_1} - a_1\right) = c_1, \quad 0 < x < \mathcal{S}, \tag{17}$$

where $c_1$ is an arbitrary constant. At steady state the forces at the interface, $x = \mathcal{S}$, are at equilibrium, giving,

$$\frac{k_1}{\eta}\left(\frac{1}{q_1} - a_1\right) = \frac{k_2}{\eta}\left(\frac{1}{q_2} - a_2\right). \tag{18}$$

Relating this mesocopic density to the microscopic density for the tissue we have

$$q_1 = \frac{N_1}{\mathcal{S}}, \; q_2 = \frac{N_2}{L - \mathcal{S}}. \tag{19}$$

Substituting Equation (19) into Equation (18) and rearranging gives

$$\mathcal{S} = \frac{\dfrac{k_1 a_1}{k_2} + \dfrac{L}{N_2} - a_2}{\dfrac{k_1}{k_2 N_1} + \dfrac{1}{N_2}}. \tag{20}$$

### S4 Breast cancer detection case study: model implementation

Here we present our method to obtain a user specified steady state cell stiffness histogram consistent with the initial $k_i$ by choosing the initial condition for the resting spring length, $a_i$.

#### S4.1 Choosing the resting spring length to choose the steady state

The possible final steady state spring stiffness histogram distributions must be consistent with the initial $k_i$ as spring properties are constant along cell trajectories. For illustrative purposes we choose the steady state spring stiffness histogram distribution we wish to obtain as guided by experimental results [5]. With this choice we can read off the histogram frequencies that describe the coverage of the each histogram interval at steady state. For simplicity, we assume that cells with $k_i$ in the same histogram interval are of equal length at steady state. Then the length of each cell in a histogram interval is given by the total length of cells in the histogram interval divided by the number of cells in the interval. As we have an initial ordering of the cells and each cell length at steady state we can now readily determine the steady state cell boundary positions. Returning to Equation (1), with a single spring per cell, at steady state and the fixed boundary conditions $x_1 = 0, x_{N+1} = L$, we now know every $x_i$ and $k_i$ and we can solve this system of $N + 1$ nonlinear equations to find each $a_i$, using fsolve in MATLAB [1]. A simulation is then initiated with these $a_i$. This simulation reaches the steady state spring stiffness histogram distribution we chose to obtain.

### S5 Supplementary Figures

In this section we present the supplementary figures which are referenced in the main text.

*Heterogeneous $k$ and homogeneous $a$.* Even though $a$ is homogeneous, the resting cell length is still a key factor as it determines whether the system is, on average, in extension or in compression. This is determined by comparing the resting spring length, $a_i^{(\nu)}$, with the critical value, $a_{\text{crit}} = L/(mN)$, which is the average length of a spring. In Figure 3 we consider $a < a_{\text{crit}}$ so that on average cells are in extension. We present other cases, where $a = a_{\text{crit}}$ and $a > a_{\text{crit}}$, in Supplementary Figures S3 and S4. We choose the initial cell stiffness distribution to be $k_0(x) = 1 + 0.1[x - (L/2)]^2$. Figure S3, depicts how the system relaxes to a uniform density distribution, due to the combination of $a = a_{\text{crit}}$ and the cell stiffness heterogeneity, as the velocity field $u$ tends to zero.

It is notable that even for low $m$ we have excellent agreement between the discrete density and the continuum density especially at the centre of each spring. However, agreement at the spring boundaries does not hold as well for low $m$.

*Heterogeneous $a$.* We see similar discrete-continuum agreement when we consider examples with homogeneous $k$ and heterogeneous $a$, and heterogeneous $k$ and heterogeneous $a$, see Supplementary Figures S5 and S6, respectively. We observe higher cell density in regions of lower $a$ which agrees with the steady state solution to the coarse-grained model, Equation (3.2) in the main paper.

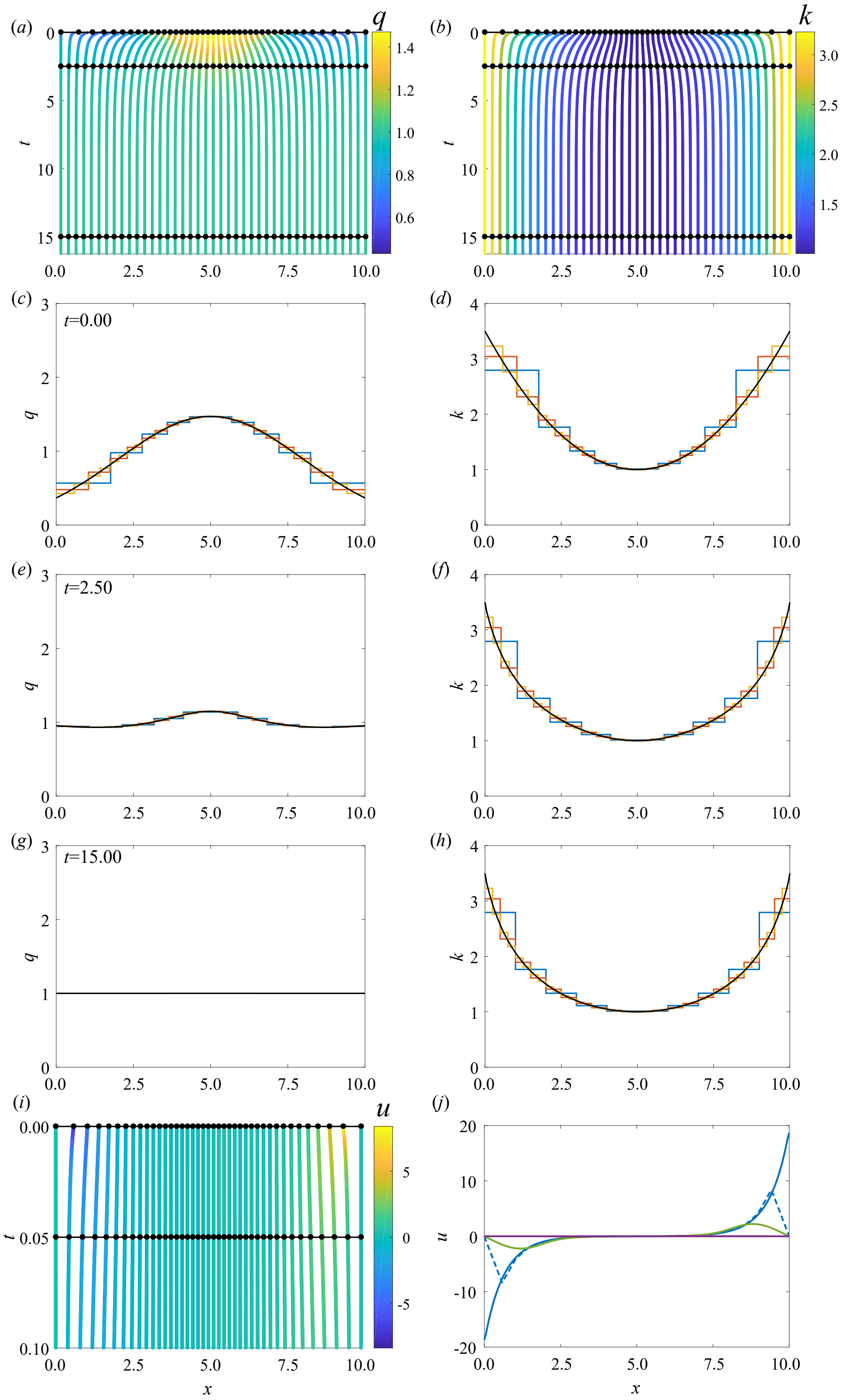

Figure S3: Results for heterogeneous cell stiffness and homogeneous cell spring length for cells in extension, with $N = 10$, $k_0(x) = 1 + 0.1(x-5)^2$, and $a_0(x) = 1$. $(a,b)$ Characteristic diagram for spring boundary position evolution for $0.00 \leq t \leq 16.25$ with $m = 4$ so that every fourth trajectory represents a cell boundary. Colour denotes $(a)$ cell density, $(b)$ cell stiffness. In $(a,b)$ black lines and dots represent times for snapshots in $(c\text{-}h)$. $(c,e,g)$ Cell density snapshots at $t = 0.00, 2.50, 15.00$. $(d,f,h)$ Cell stiffness snapshots at $t = 0.00, 2.50, 15.00$. In $(c\text{-}h)$ lines display results from $m = 1$ (blue), 2 (red), 4 (yellow), and continuum system (black). $(i)$ Characteristic diagram for spring boundary position evolution for $0.00 \leq t \leq 0.10$. Colour denotes velocity. $(j)$ Velocity snapshots at $t = 0.00$ (blue), 0.05 (green), 2.50 (yellow), 15.00 (magenta). Dashed/solid line represent solutions from discrete model with $m = 4$ and continuum model, respectively.

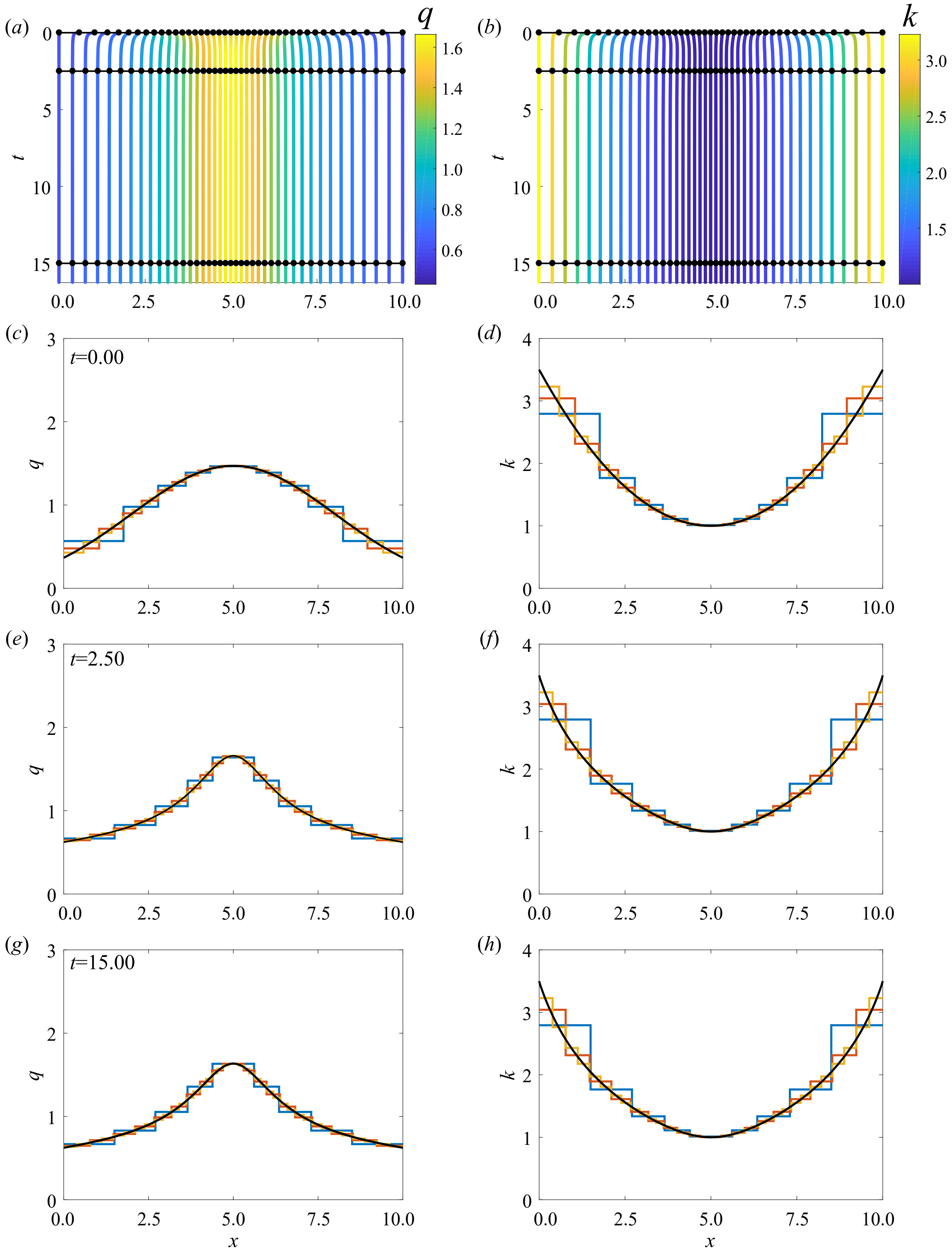

Figure S4: Results for heterogeneous cell stiffness and homogeneous resting cell length, for cells on average in compression, with $N = 10$, $k_0(x) = 1 + 0.1(x-5)^2$, and $a_0(x) = 2$. (*a*,*b*) Characteristic diagram for spring boundary position evolution for $0.00 \leq t \leq 16.25$ with $m = 4$ so that every fourth trajectory represents a cell boundary. Colour denotes (*a*) cell density, (*b*) cell stiffness. In (*a*,*b*) black lines and dots represent times for snapshots in (*c*-*h*). (*c*,*e*,*g*) Cell density snapshots at $t = 0.00, 2.50, 15.00$. (*d*,*f*,*h*) Cell stiffness snapshots at $t = 0.00, 2.50, 15.00$. In (*c*-*h*) lines display results from $m = 1$ (blue), 2 (red), 4 (yellow), and continuum system (black).

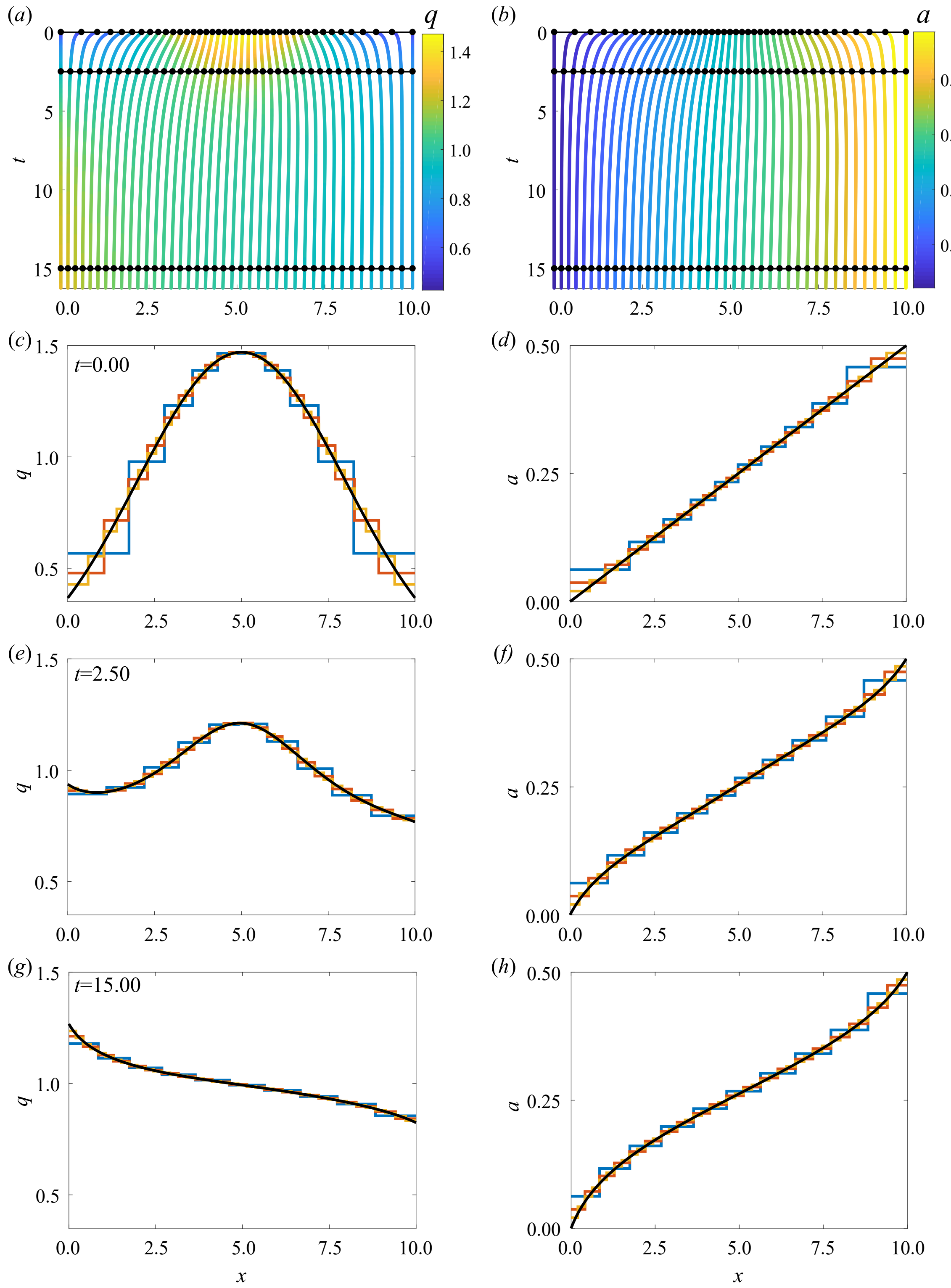

Figure S5: Results for homogeneous $k$ and heterogeneous $a$ for cells on average not in extension or compression, with $N = 10$, $k_0(x) = 1$, and $a_0(x) = 0.05x$. ($a$,$b$) Characteristic diagram for spring boundary position evolution for $0.00 \leq t \leq 16.25$, with $m = 4$ so that every fourth trajectory represents a cell boundary. Colour denotes ($a$) cell density, ($b$) resting cell length. In ($a$,$b$) black lines and dots represent times for snapshots in ($c$-$h$). ($c$,$e$,$g$) Cell density snapshots at $t = 0.00, 2.50, 15.00$. ($d$,$f$,$h$) Resting cell length snapshots at $t = 0.00, 2.50, 15.00$. In ($c$-$h$) lines display results from $N = 10$ with $m = 1$ (blue), 2 (red), and continuum system (black).

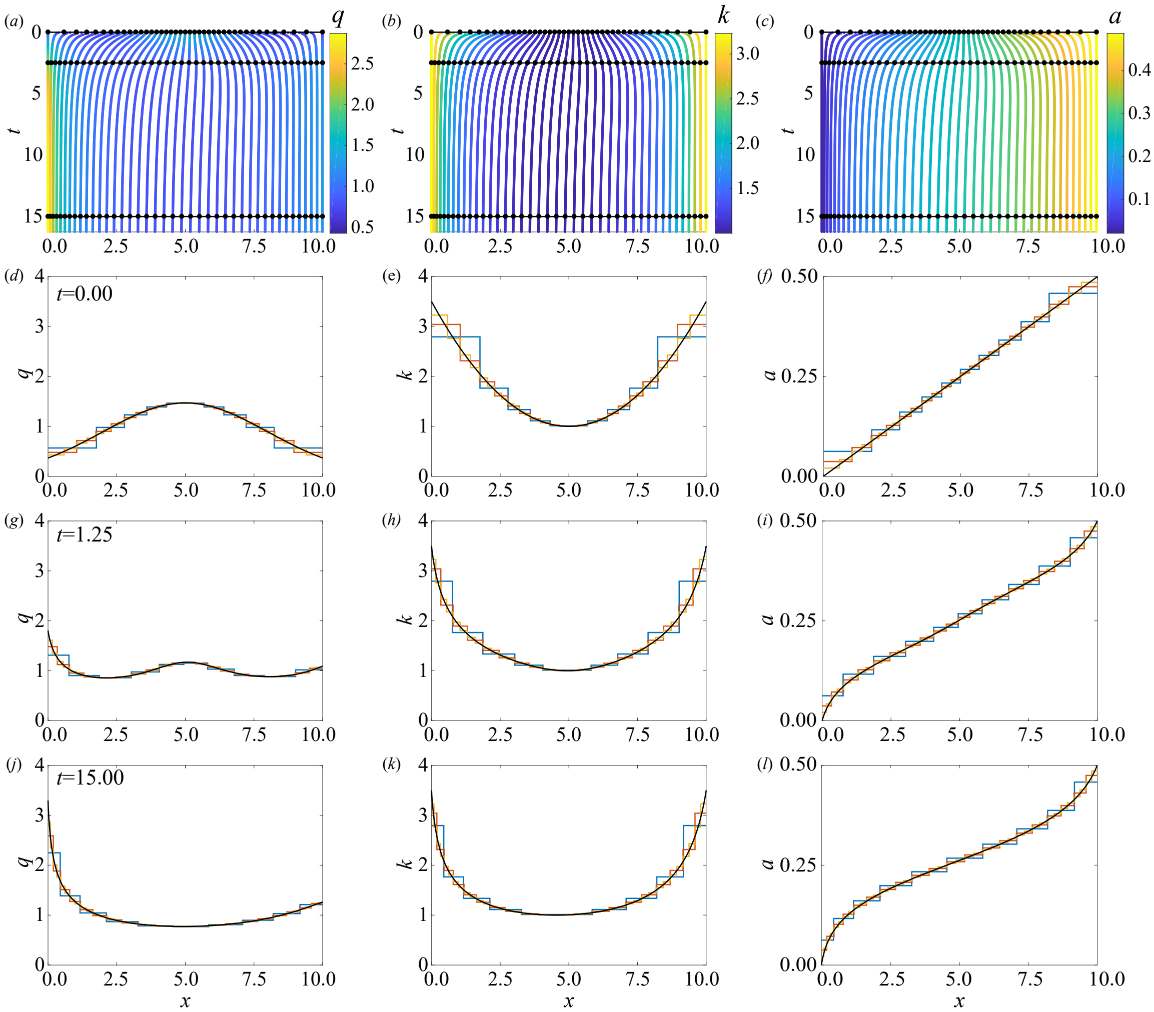

Figure S6: Results for heterogeneous cell stiffness and heterogeneous resting cell length, with $N = 10$, $k_0(x) = 1 + 0.1(x-5)^2$ and $a_0(x) = 0.05x$. (*a,b,c*) Characteristic diagram for spring boundary position evolution for $0.00 \le t \le 16.25$ with $m = 4$ so that every fourth trajectory represents a cell boundary. Colour denotes (*a*) cell density, (*b*) cell stiffness, (*c*) resting cell length. In (*a,b,c*) black lines and dots represent times for snapshots in (*d*-*l*). (*d,g,j*) Cell density snapshots at $t = 0.00, 1.25, 15.00$. (*e,h,k*) Cell stiffness snapshots at $t = 0.00, 1.25, 15.00$. (*f,i,l*) Resting cell length snapshots at $t = 0.00, 1.25, 15.00$. In (*d*-*l*) lines display results from $m = 1$ (blue), 2 (red), 4 (yellow), and continuum system (black).

### References

[1] Mathworks. 2019 *Optimization Toolbox: User's Guide (R2019a)*. Retrieved March 27, 2019 from https://au.mathworks.com/help/pdf_doc/optim/optim_tb.pdf

[2] Shampine LF, Reichelt MW. 1997 The MATLAB ODE Suite. *SIAM Journal on Scientific Computing.* **18**. 1-22. (doi:10.1137/S1064827594276424)

[3] Zheng C, Bennett G. 2002. *Applied Contaminant Transport Modeling.* New York, NY: Wiley-Blackwell.

[4] Crank J. 1975 *The Mathematics of Diffusion.* 2nd ed. Oxford, UK: Oxford University Press.

[5] Plodinec M, Loparic M, Monnier CA, Obermann EC, Zanetti-Dallenbach R, Oertle P, Hyotyla JT, Aebi U, Bentires-Alj M, Lim RYH, et al. 2012 The nanomechanical signature of breast cancer. *Nat. Nanotechnol.* **7**. 757-764. (doi:10.1038/nnano.2012.167)
